# A comparative analysis of memory B cell and antibody responses against *Plasmodium falciparum* merozoite surface protein 1 in children and adults from Uganda

**DOI:** 10.1101/2021.11.04.467302

**Authors:** S. Jake Gonzales, Kathleen N. Clarke, Gayani Batugedara, Ashley E. Braddom, Rolando Garza, Raphael A. Reyes, Isaac Ssewanyana, Kendra C. Garrison, Gregory C. Ippolito, Bryan Greenhouse, Sebastiaan Bol, Evelien M. Bunnik

## Abstract

Memory B cells (MBCs) and plasma antibodies against *Plasmodium falciparum* merozoite antigens are important components of the protective immune response against malaria. To gain understanding of how responses against *P. falciparum* develop in these two arms of the humoral immune system, we evaluated MBC and antibody responses against the most abundant merozoite antigen, merozoite surface protein 1 (MSP1), in individuals from a region in Uganda with high *P. falciparum* transmission. Our results showed that MSP1-specific B cells in adults with immunological protection against malaria were predominantly IgG^+^ classical MBCs, while children with incomplete protection mainly harbored IgM^+^ MSP1-specific classical MBCs. In contrast, anti-MSP1 plasma IgM reactivity was minimal in both children and adults. Instead, both groups showed high plasma IgG reactivity against MSP1 and whole merozoites, with broadening of the response against non-3D7 strains in adults. The antibodies encoded by MSP1-specific IgG^+^ MBCs carried high levels of amino acid substitutions and recognized relatively conserved epitopes on the highly variable MSP1 protein. Proteomics analysis of MSP1_19_-specific IgG in plasma of an adult revealed a limited repertoire of anti-MSP1 antibodies, most of which were IgG_1_ or IgG_3_. Similar to MSP1- specific MBCs, anti-MSP1 IgGs had relatively high levels of amino acid substitutions and their sequences were predominantly found in classical MBCs, not atypical MBCs. Collectively, these results showed evolution of the MSP1-specific humoral immune response with cumulative *P. falciparum* exposure, with a shift from IgM^+^ to IgG^+^ B cell memory, diversification of B cells from germline, and stronger recognition of MSP1 variants by the plasma IgG repertoire.

## Introduction

Malaria, caused by the parasite *Plasmodium falciparum*, is responsible for approximately half a million deaths every year, of which two-thirds occur in children under the age of five (1). A much larger number of people experience non-fatal malaria, amounting to an estimated 230 million cases of disease annually. Although the mortality rate for malaria has slowly but consistently declined over the past two decades, the decrease in malaria incidence has plateaued in the past five years. Current interventions are thus insufficient for malaria elimination and novel tools, such as a highly efficacious malaria vaccine, are urgently needed in the fight against this devastating disease. RTS,S, the only malaria vaccine that has elicited protection against (severe) malaria in a phase III clinical trial, had an efficacy against clinical malaria of 30 – 40% in infants and young children (2). Vaccine efficacy was high shortly after vaccination but declined rapidly, and was lower against parasites that were genetically different from strain 3D7 that the subunit vaccine RTS,S was based on (3). Compared to vaccination, repeated natural *P. falciparum* infections eventually elicit superior immunity, consisting of relatively long-lived antibody responses (∼2 – 4 years) with cross-strain reactivity (4, 5). This naturally acquired humoral immunity against malaria is associated with the presence of circulating immunoglobulin G (IgG) against *Plasmodium* blood stage antigens (6–13). Recently, immunoglobulin M (IgM) has received increased attention as IgM responses were also shown to correlate with protection and were able to inhibit parasite growth *in vitro* (14–16). A complete understanding of the development of humoral immune responses against *P. falciparum* blood stage antigens over time will help with the development of a more effective and longer lasting vaccine than RTS,S.

Effective and long-lasting humoral immune responses against pathogens consist of long-lived memory B cells (MBCs) in the circulation and plasma cells that reside in the bone marrow and secrete large amounts of antibody into the circulation (reviewed in (17)). In the case of *P. falciparum*, long-lived humoral immunity develops over the course of years of repetitive infections, leaving young children largely susceptible to disease. With cumulative *P. falciparum* exposure, the quality of B cell responses gradually improves. Both MBCs and plasma cells generated in response to *P. falciparum* infection are short-lived in young children but gradually increase in longevity in adolescents and adults (4, 18, 19). Over time, antibody responses also recognize a larger number of parasite antigens (18). In addition, recurrent infections drive the generation of antibodies capable of mediating cross-strain immunity, which is strongly associated with protection (5). Most studies into the development of antibody responses against *P. falciparum* have been performed using serum. However, this prevents investigating the molecular characteristics of *P. falciparum*-specific antibodies. In addition, little is known about the connection between the MBC and plasma cell compartments. This information will be useful for understanding how durable immunity against malaria develops and may enable us to harness lessons from naturally acquired immunity for improved vaccine design.

In this study, we set out to investigate differences in humoral immune responses in between children with incomplete protection against malaria and adults who have developed strong immunological protection to better understand how MBC and antibody responses develop over the course of life-long *P. falciparum* exposure. To do this, we compared the antibody and MBC response against *P. falciparum* antigen merozoite surface protein 1 (MSP1) between children and adults living in a region of high *P. falciparum* transmission in Uganda. MSP1 is a large, polymorphic, and highly immunogenic protein expressed ubiquitously on the surface of the parasite during the late schizont and merozoite stages (20, 21). MSP1 has long been considered a vaccine target since antibody responses against this protein have been associated with protection (6, 20, 22), although results from MSP1-based vaccine trials have been disappointing (23, 24). However, since MSP1 has high antigenic heterogeneity across *P. falciparum* strains, we considered it an excellent model antigen to assess antibody cross-strain reactivity elicited by natural infection. We studied the isotype of MSP1- specific MBCs, sequence characteristics of monoclonal antibodies isolated from IgM^+^ and IgG^+^ MBCs, and cross-strain reactivity against MSP1 variants in both plasma antibodies and B cells. In addition, we analyzed the anti-MSP1 plasma IgG repertoire and investigated characteristics of antibodies that were found in both the MBC and plasma IgG compartment in more detail.

## Results

### MSP1-specific B cells have a classical phenotype and are enriched for IgG in adults

To study differences in naturally acquired B cell responses against MSP1 between partially immune and immune individuals, we selected four children and four adults residing in Tororo, Uganda, a region with high malaria transmission intensity year-round (**Supplementary table 1**) (25). These individuals were participants of cohort studies, with the exception of two Ugandan adults who were anonymous blood donors with high levels of plasma antibodies against various malaria antigens, suggesting frequent exposure to *P. falciparum* (**Supplementary figure 1**). Cryopreserved PBMCs were used to first isolate bulk B cells, followed by staining with fluorescently labeled tetramers of full-length MSP1 from *P. falciparum* strain 3D7 (MSP1_3D7_) and decoy tetramers for analysis by flow cytometry, a strategy developed and used by others (26, 27) (**Figure 1A**). In both children and adults, MSP1-specific B cells (defined as CD19^+^CD20^+^MSP1^+^decoy^-^) were predominantly found among CD21^+^CD27^+^ classical memory B cells (MBCs) (**Figure 1B**). In addition, in both groups, very few MSP1- specific B cells were found among CD21^-^CD27^-^ atypical MBCs (**Figure 1B**). To further define the phenotype of MSP1-specific B cells, we determined the isotype usage of MSP1-specific classical MBCs. Children showed much larger percentages of IgM^+^ (median, ∼70%) than IgG^+^ (median, ∼10%) classical MBCs in the total repertoire, which was reflected in the percentage of IgM^+^ and IgG^+^ MSP1-specific classical MBCs (**Figure 1C**). In contrast, IgM^+^ and IgG^+^ classical MBCs were equally abundant in adults in the total repertoire, while IgM^+^ MSP1-specific classical MBCs were depleted and IgG^+^ MSP1-specific classical MBCs were slightly enriched (**Figure 1C**). The ratio of IgG^+^ to IgM^+^ MSP1-specific classical MBCs was higher in all adults as compared to the children (**Figure 1D**). Collectively, these results suggest that the B cell response against MSP1 is skewed to IgM^+^ classical MBCs in children, while it is dominated by IgG^+^ classical MBCs in adults. These results are in line with a recent report that showed an increased frequency of IgM usage among *P. falciparum*-specific B cells in young children as compared to older children and adults in Mali, where malaria transmission is highly seasonal (16), suggesting that this is a general feature of the B cell response to *P. falciparum* in children in malaria-endemic regions.

**Figure 1:**
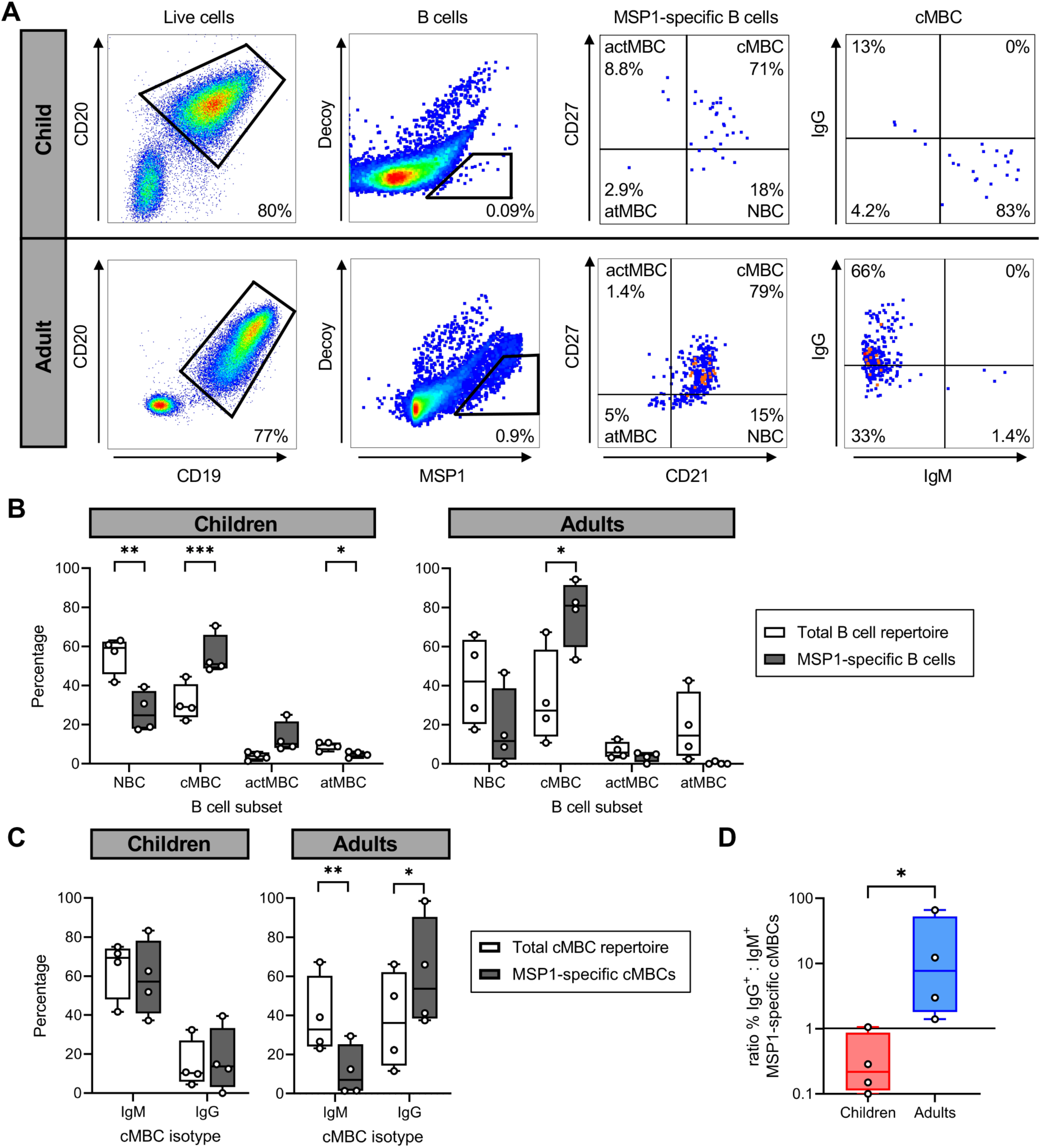
The isotype of MSP1-specific B cells in malaria-experienced children and adults. **A)** Representative flow cytometry gating for the sorting and analysis of MSP1-specific B cells. The four major B cell subsets are defined as follows: naïve B cells (NBC), CD21^+^ CD27^-^; classical memory B cells (cMBC), CD21^+^ CD27^+^; activated memory B cells (actMBC), CD21^-^ CD27^+^; and atypical memory B cells (atMBC), CD21^-^ CD27^-^. Data are shown as pseudocolor plots, in which overlapping cells in the plots showing MSP1-specific B cells and cMBCs are shown in orange and red. **B)** Relative abundance of major B cell subsets among the total B cell repertoire and among MSP1- specific B cells in partially immune children (n = 4) and immune adults (n = 4). **C)** The percentage of IgM^+^ and IgG^+^ B cells among the total repertoire of cMBCs and among MSP1-specific cMBCs in malaria-experienced children (n = 4) and adults (n = 4). In panels B and C, differences were tested for statistical significance using a paired Student’s t-test, even though a non-parametric test would be more appropriate given the small sample size. However, the paired non-parametric alternative (Wilcoxon signed-rank test) does not return a P value lower than 0.13 when using groups of 4. **D)** The ratio of the percentage of IgG^+^ over IgM^+^ MSP1-specific cMBCs in malaria-experienced children and adults. A data point with value 0 was plotted at 0.1 for visualization purposes. The difference between groups was tested for statistical significance using a Mann Whitney test. *** P < 0.001; ** P < 0.01; * P < 0.05

### Recombinant antibodies from isolated memory B cells are MSP1-specific and inhibit parasite growth

To confirm antigen-specificity of MSP1-specific B cells, all IgM^+^ and IgG^+^ MSP1-specific B cells were single-cell-sorted into 96-well culture plates containing CD40L-expressing feeder cells, cytokines, and other stimuli that collectively promote B cell survival, expansion, and differentiation into antibody-secreting cells. This allowed us to screen B cell clones for antigen-specificity prior to cloning and expression of recombinant mAbs. B cell supernatants were tested for the presence of anti-MSP1 antibodies by a *P. falciparum* strain 3D7 merozoite ELISA or Luminex assay using recombinant MSP1_3D7_. These assays were implemented into our analysis pipeline only in later experiments and these data are therefore missing for samples analyzed earlier. The variable regions of antibodies with confirmed reactivity to merozoites or recombinant MSP1 were cloned into linear expression cassettes, expressed as recombinant IgG_1_, and against tested for MSP1 reactivity (**Figure 2A**). As was reported in a recently published article (15), most anti-MSP1 IgM antibodies lost their reactivity to MSP1 when expressed as IgG_1_, which is likely a result of the lower avidity of monomeric IgG as compared to pentameric IgM. We therefore cloned a multimerization domain into the C-terminus of the IgG_1_ heavy chain to express IgM-derived antibodies as pentameric IgG. In contrast to results reported by Thouvenel *et al*. (15), we were unable to rescue the MSP1-reactivity of IgM-derived variable regions by expression as pentameric IgG (**Figure 2A**). The only pentameric IgG with reactivity to MSP1 was also reactive when expressed in monomeric form. To further confirm the specificity and functionality of the isolated anti- MSP1 mAbs, we performed immunofluorescence assays on segmented schizonts for three randomly selected IgG mAbs, showing the expected surface staining of merozoites (**Figure 2B**). We also performed growth-inhibition assays for six IgG mAbs that were selected based on high expression levels in culture. At a concentration of 200 µg/ml, these six mAbs showed on average 51% (range, 17% – 65%) inhibition of *P. falciparum* growth as compared to a negative control, demonstrating their ability to functionally inhibit the parasite (**Figure 2C**).

**Figure 2:**
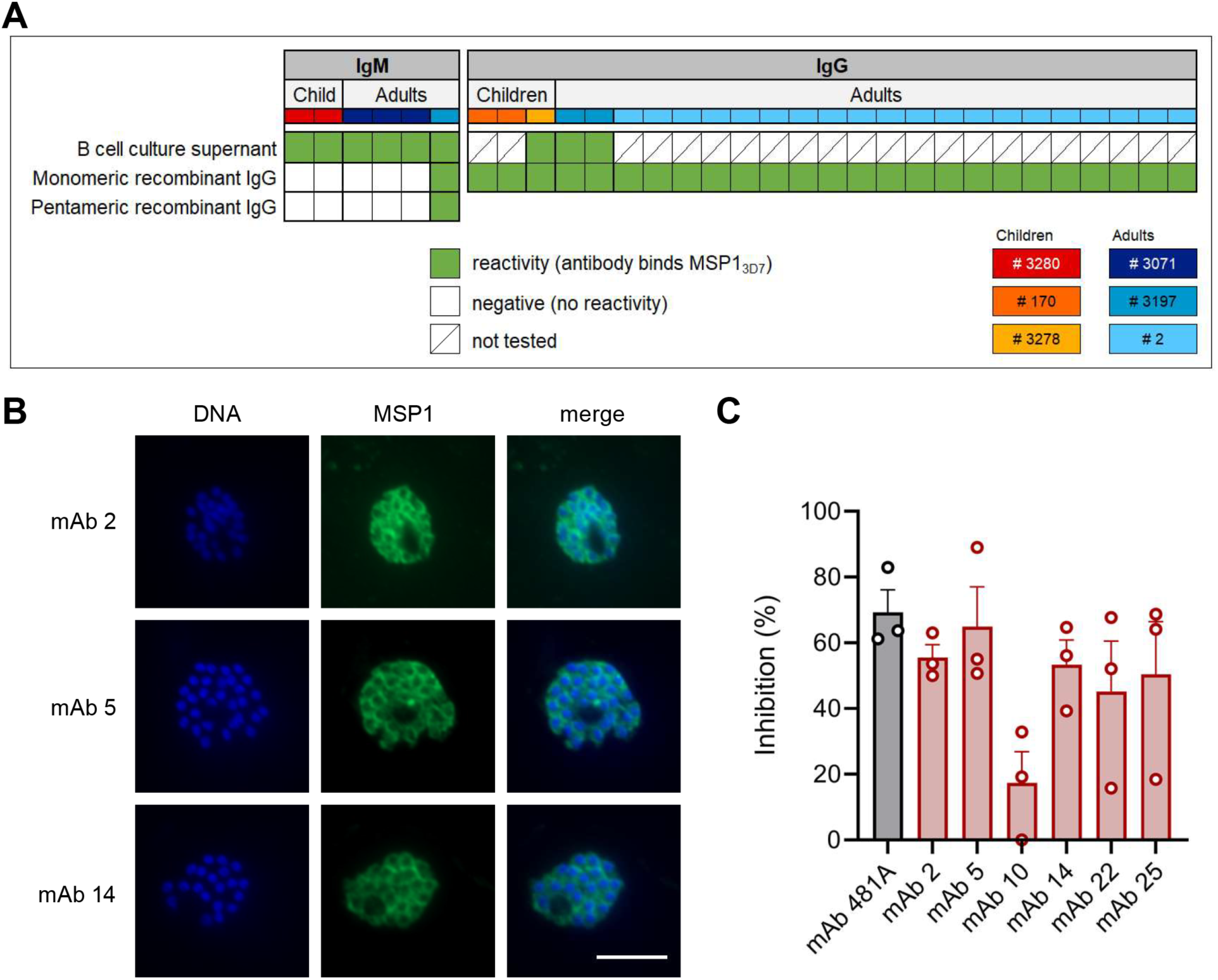
MSP1_3D7_ reactivity and growth inhibitory activity of anti-MSP1 monoclonal antibodies isolated from memory B cells. **A)** MSP1_3D7_ reactivity of supernatants of clonal MBC cultures or mAbs obtained from MBCs (recombinant IgG) determined by ELISA or Luminex assay. The donor from whom mAbs were isolated is color-coded in the top row. Recombinant mAbs that were derived from IgM^+^ MBCs were tested as both monomeric and pentaimeric recombinant IgG. **B)** Immunofluorescence imaging of segmented schizonts using three select mAbs isolated from MBCs of adult donor 2. At this stage, parasites consist of 16 – 32 merozoites, each with its own nucleus, and express MSP1 on the parasite cell surface. Representative images of two experiments are shown. Scale bar is 5 μm. **C)** *P. falciparum* growth inhibition by six select mAbs isolated from adult donor 2. Inhibition was calculated relative to a negative control. Results shown are the average + SEM from three independent experiments. mAb 481A served as a positive control.

### MSP1-specific IgG^+^ MBC lineages from a malaria-experienced adult are highly diversified and expanded

Next, we set out to study the molecular characteristics of antibodies encoded by MSP1- specific MBCs. In total, we isolated 6 IgM and 25 IgG mAbs with confirmed MSP1 reactivity from 6 donors. Full-length heavy and light chain variable regions were obtained and analyzed using IMGT/V-QUEST (28) to determine the level of amino acid substitutions in the V gene segments of the variable regions. MSP1-specific IgM^+^ MBCs from adults and children carried few amino acid substitutions in both the heavy and light chain V gene segments (average, <1% amino acid changes), while many of the IgG^+^ MBCs were highly mutated (>15% amino acid changes; **Figure 3A,B**). None of the IgM^+^ MBCs were clonally related to each other, defined as sequences using the same heavy chain V and J gene and having a highly similar heavy chain complementarity determining region 3 (HCDR3; ≥85% amino acid similarity). In contrast, several clonally expanded IgG^+^ MBC lineages were identified in adult donor 2, from whom most mAbs were derived (**Figure 3A,B**). B cells belonging to expanded lineages were among those that were the most diversified from the germline antibody sequences, particularly in the heavy chain V gene segment. Finally, although there was no statistically significant difference in the length of the HCDR3 between IgM and IgG sequences, several IgG sequences harbored long HCDR3s (≥20 amino acids), while IgM sequences had average or relatively short HCDR3s (**Figure 3C**).

**Figure 3:**
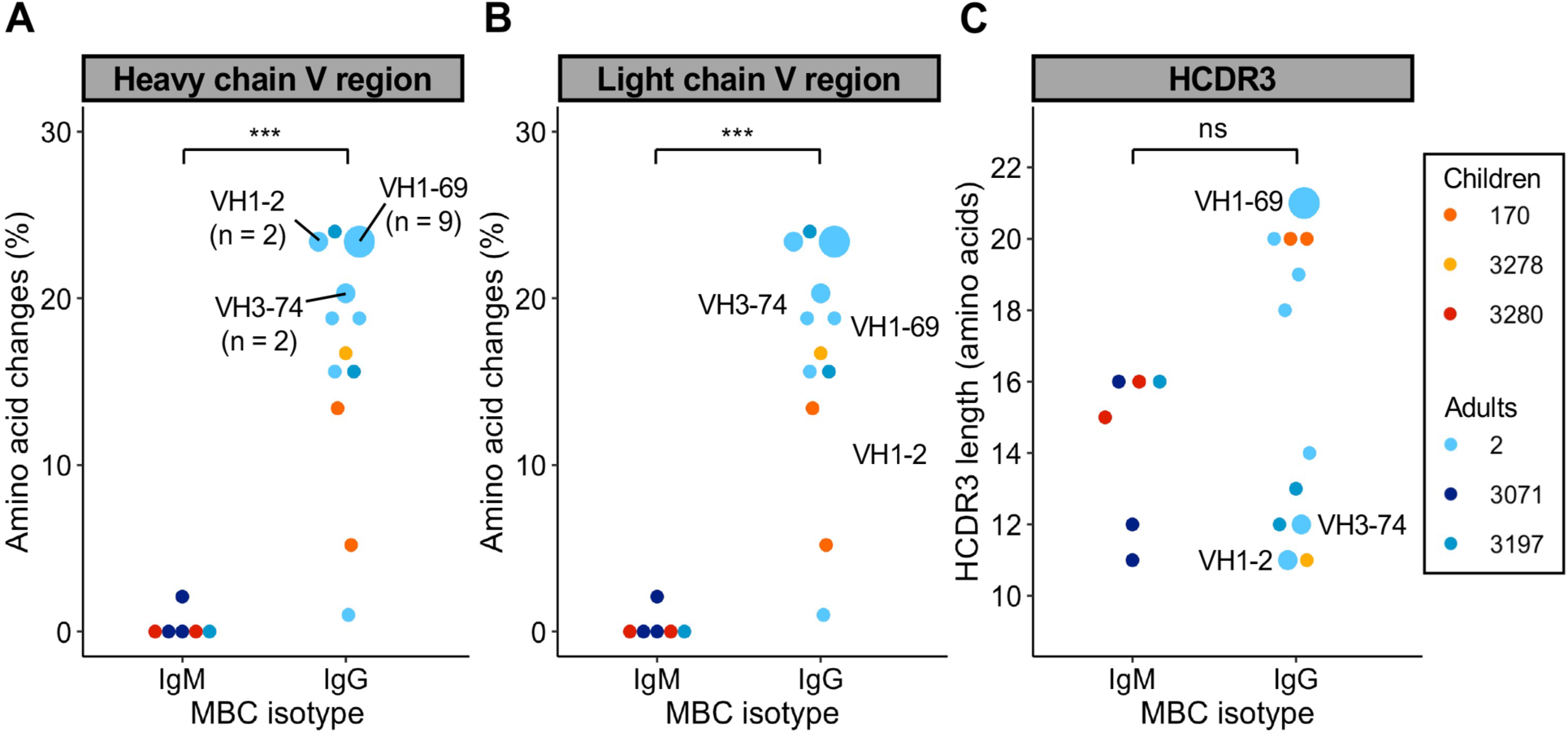
Sequence analysis of the variable heavy and light chains of MSP1-specific memory B cells. **A-B)** Percentage of amino acid changes in the V gene segments of the heavy chain (A) and light chain (B) variable regions. **C)** Length of HCDR3s. In all graphs, sequences from expanded clonal lineages were grouped. The VH gene usage and size of expanded clonal lineages are indicated. Differences between IgM^+^ (n = 6) and IgG^+^ (n = 13 for heavy chains / n = 14 for light chains) MBC lineages were evaluated for statistical significance using a Mann Whitney test.

### Anti-MSP1 plasma antibody responses are dominated by IgG in both children and adults

A study of B cell responses upon influenza virus vaccination reported that the largest clonal B cell families were directed against the most conserved epitopes (29). Based on these observations, we hypothesized that clonally expanded anti-MSP1 IgG^+^ MBC lineages would have cross-strain reactivity, whereas non-expanded IgG^+^ MBC lineages would be strain-specific. To test this hypothesis, we developed a Luminex assay using beads coated with full-length MSP1 variants from the geographically and genetically distinct *P. falciparum* strains 3D7 (African origin), Dd2 (South-east Asian origin), and HB3 (Central American origin). 21 out of 22 mAbs (95%) tested in this assay bound to MSP1_3D7_ (**Figure 4A**). This high reactivity with MSP1_3D7_ was expected since MSP1_3D7_ was used for the isolation of MSP1-specific B cells. It is therefore unclear why one mAb bound MSP1_Dd2_ but not MSP1_3D7_. All other mAbs showed reactivity with at least one other MSP1 variant, and the majority showed reactivity with both MSP1_Dd2_ and MSP1_HB3_, including all mAbs from the largest clonal lineage (**Figure 4A**). These results suggest that the majority mAbs recognized relatively conserved epitopes, irrespective of the level of clonal expansion of the B cell lineage. Unfortunately, we were unable to determine reactivity of mAbs isolated from IgM^+^ MBCs against the three MSP1 variants from different *P. falciparum* strains, because of the loss of antigen reactivity when these mAbs were expressed as IgG. In addition, most mAbs were derived from a single donor, limiting the conclusions we could draw from this experiment.

**Figure 4:**
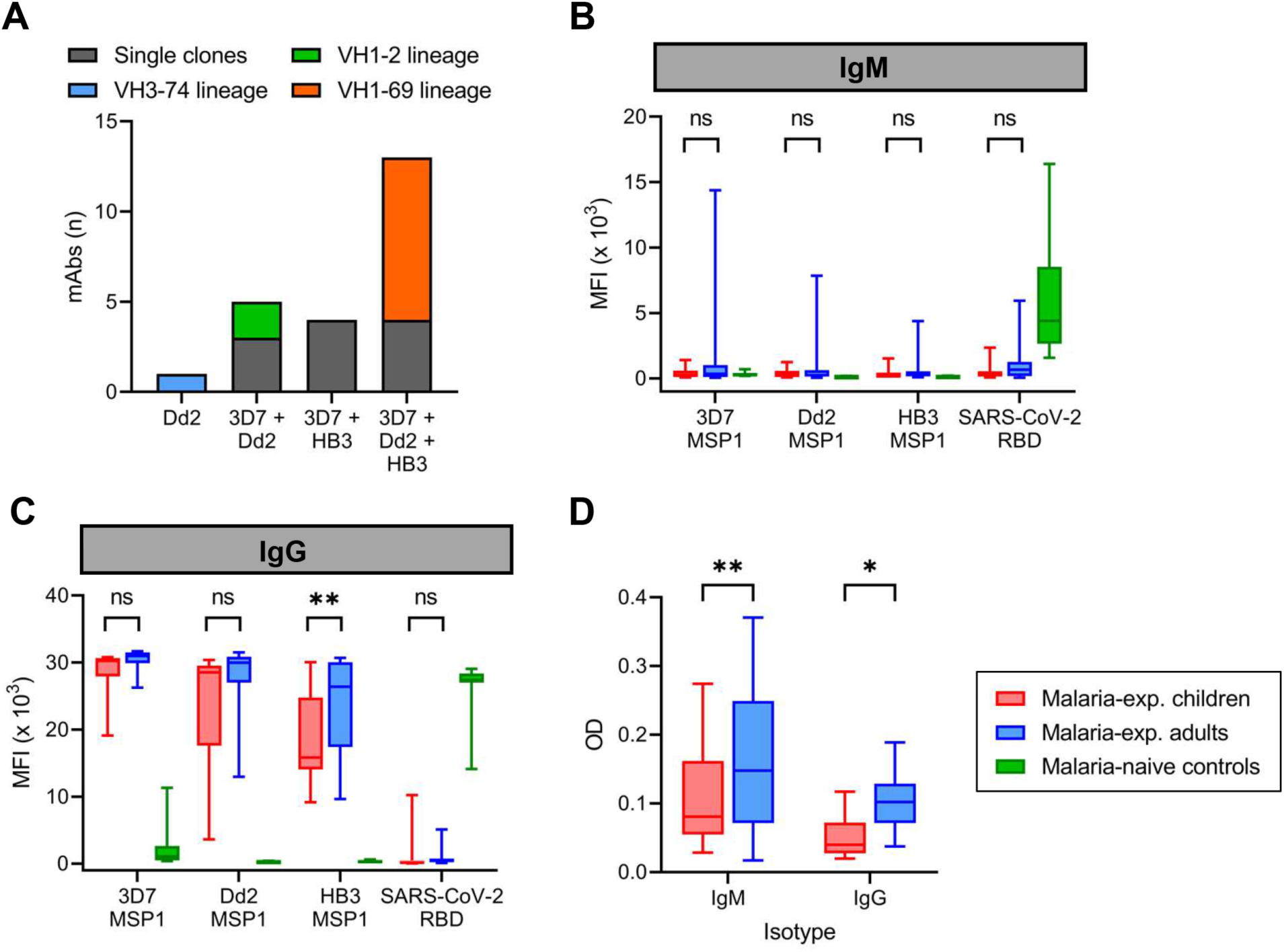
Reactivity of monoclonal antibodies and plasma IgM and IgG against different strains of *P. falciparum.* **A)** The number of anti-MSP1 recombinant monoclonal antibodies (mAbs) with reactivity against recombinant MSP1 of three different *P. falciparum* strains in a Luminex assay. Each mAb is shown as a stacked block and is color-coded based on whether it was a single clone or derived from an expanded clonal B cell lineages. The VH3-74 lineage consisted of two MBCs, of which we were unable to retrieve the light chain for the second mAb. For this reason, only a single mAb of this lineage was tested here. **B-C)** Reactivity of plasma IgM (B) and IgG (C) from malaria-experienced children (n = 18) and adults (n = 24), and malaria-naïve recovered US COVID-19 patients (control; n = 10) against recombinant MSP1 of *P. falciparum* strains 3D7, Dd2, and HB3, as well as SARS-CoV-2 receptor binding domain (RBD), as determined by Luminex assay. Values shown are mean fluorescence intensity (MFI). **D)** Reactivity of plasma IgM and IgG from malaria-experienced children (n = 18) and adults (n = 24) against whole merozoites by ELISA. Values shown are optical densities after subtraction of background values obtained using pooled unexposed plasma from US donors. Data shown in panels B – D were tested for statistical significance using a two-way repeated measures ANOVA, followed by comparisons between children and adults using Šídák’s post-hoc test, corrected for multiple comparisons. ** P < 0.01; * P < 0.05

To expand on the results of cross-strain reactivity of mAb against MSP1 and to determine whether the MSP1-specific MBC isotype differences we observed between children and adults are reflected in the plasma antibody response, we tested plasma samples from 24 adults and 18 children for IgM and IgG reactivity against MSP1 from the three *P. falciparum* strains (**Supplementary table 2**). This experimental design provides a measurement of the overall antibody reactivity of the plasma but does not discriminate between antibodies with cross-strain reactivity or a combination of multiple strain-specific antibodies. However, it will reveal whether anti-MSP1 antibody responses broaden with cumulative *P. falciparum* exposure. As a control, we included plasma samples from recovered US COVID-19 patients who showed IgM and IgG reactivity against the SARS-CoV-2 receptor binding domain, but not against the three *P. falciparum* MSP1 variants. In both children and adults, anti-MSP1 IgM reactivity in the plasma was very low, although four of the 24 adult samples contained anti-MSP1 IgM against at least one MSP1 variant antigen (**Figure 4B**). In contrast, all samples showed strong IgG reactivity against all MSP1 variants (**Figure 4C**). Adults had higher plasma IgG reactivity against MSP1_HB3_ than children. The average reactivity of adult samples against MSP1_Dd2_ was also higher but not statistically significantly different from that in children (P = 0.06). These results suggest that while children already have plasma IgG against different variants of MSP1, this response continues to broaden with age.

To determine whether the differential IgM and IgG reactivity of plasma is specific for anti-MSP1 antibodies or can be extrapolated to anti-merozoite antibody responses in general, we also tested reactivity of plasma IgM and IgG to whole merozoites from *P. falciparum* strain 3D7. In this assay, we observed anti-IgM and anti-IgG reactivity in both groups, with adults showing higher reactivity for both IgM and IgG as compared to children (**Figure 4D**). IgM and IgG anti-merozoite reactivity were not directly compared since the values measured could have been influenced by the binding affinity of the secondary antibody. Collectively, these results suggest that in contrast to our observation that the MBC response in children is enriched for IgM^+^ MBCs, the plasma antibody response against MSP1 in both children and adults is dominated by IgG. However, IgM responses against other merozoite antigens may be better developed, in particular in adults.

### The anti-MSP1 plasma IgG repertoire has limited diversity, high levels of amino acid substitutions, and mainly overlaps with sequences found in classical memory B cells

To further explore the connection between the plasma cell and MBC compartments of the humoral immune response, we performed an integrative analysis of the plasma anti- MSP1 IgG and B cell receptor repertoires in adult donor 2, who was selected based on the availability of additional PBMCs and plasma. Although the results of this experiment will require confirmation in additional individuals, this is the first analysis of its kind in a malaria-experienced person and will provide valuable insight into the molecular characteristics of the anti-MSP1 plasma antibody repertoire after life-long exposure to *P. falciparum*. In a previous study, we generated B cell receptor sequencing (BCR-seq) data of antibody heavy chain variable regions of naïve B cells, classical MBCs, and atypical MBCs (30). The full BCR-seq data set and all sequences obtained from MSP1- specific MBCs were used to construct a personalized, donor-specific reference heavy chain antibody variable region sequence database. We then isolated anti-MSP1 IgG from plasma using commercially available 19 kDa C-terminal fragment of MSP1 (MSP1_19_), which limited our analysis to the most conserved domain of MSP1 (31, 32). We analyzed the anti-MSP1_19_ IgG preparation by high-resolution liquid chromatography with tandem mass spectrometry and searched the obtained spectra against the donor-specific antibody variable region database (**Figure 5A**). This step allowed us to identify the full-length antibody sequences that these short peptide spectra were derived from. Eighteen anti-MSP1_19_ IgG lineages were identified, of which four lineages made up over 75% of all plasma anti-MSP1_19_ IgG (**Figure 5B, Supplementary table 3**), suggesting that the anti-MSP1 plasma IgG repertoire is relatively limited in diversity. The 18 antibody lineages were dominated by sequences that were found among IgG_1_^+^ and IgG_3_^+^ classical MBCs, all of which had relatively high levels of amino acid substitutions (>15%) (**Figure 5B**). These results suggest that MSP1-specific B cell lineages can give rise to both plasma cells and classical memory B cells, but not atypical MBCs. Two relatively abundant antibody lineages (comprising 23.5% and 6.9% of anti-MSP1_19_ plasma IgG) overlapped with IgG^+^ classical MBC lineages that were expanded in the bulk BCR-seq data (5 and 4 clonal B cell members, respectively). Both of these lineages used IGHV1-69 and IGHJ6 and had long HCDR3 sequences of 24 and 21 amino acid residues, respectively, suggesting that these characteristics may have contributed to the selection and expansion of both MBC and plasma cell populations. Long HCDR3s were also observed among the other IgG clonal lineages (average of all lineages, 18 amino acid residues; **Supplementary table 3**), although the most abundant IgG (representing 24.7% of all anti-MSP1_19_ plasma IgG) had an HCDR3 of 10 amino acid residues. A third antibody lineage with low abundance (<0.1%) matched with an expanded B cell lineage, consisting of two clonal IgM^+^ atypical MBC members that had relatively high levels of amino acid substitutions (>10%). Except for these three expanded lineages, all other clonotypes were found as a single sequence in the bulk BCR-seq data set. These results highlight that the abundance of antibody clonotypes in the plasma is not necessarily reflected by an expansion of the corresponding MBC lineages, pointing towards different selective mechanisms that govern MBC and plasma cell development.

**Figure 5:**
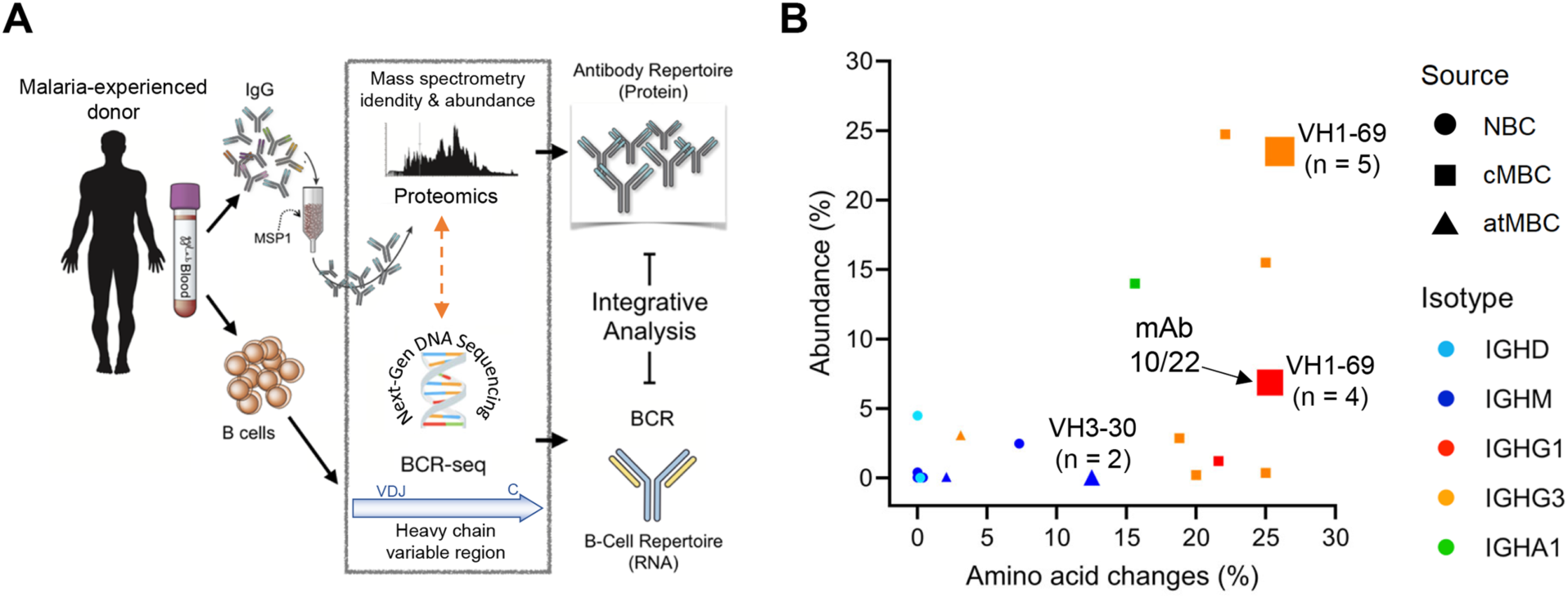
Analysis of the repertoire of anti-MSP1_19_ IgG in the plasma of a malaria-experienced adult. **A)** Schematic overview of the experimental pipeline. **B)** The percentage amino acid changes in the V gene segment of the heavy chain variable region (as determined by B cell receptor sequencing; X axis) plotted against the abundance of the corresponding IgG in plasma (Y axis). The size of the data points indicates the number of clonal sequences found in the B cell receptor sequencing (BCR-seq) data set, as a measure of the size of the clonal B cell lineage. The three lineages that were expanded in the BCR-seq data set are indicated with a number, that represents the number of clonal sequences identified by BCR-seq. The shape of the data point shows the B cell subset among which the sequence was found (NBC, naïve B cell; cMBC, classical memory B cell; atMBC, atypical memory B cell), while the color indicates the B cell isotype. The data point corresponding to mAb10 and mAb22 (isolated from MSP1-specific B cells) is indicated with an arrow.

### Analysis of an expanded B cell lineage detected in both plasma IgG and memory B cells

To further analyze antibody characteristics that may influence their selection and expansion among plasma cells and MBCs, we compared the anti-MSP1 IgG sequences detected in plasma with those derived from MSP1-specific MBCs. Only the largest MSP1-specific mAb lineage was detected among the anti-MSP1_19_ antibody lineages identified in plasma, potentially because other MSP1-specific mAbs target epitopes in other parts of the MSP1 protein. Two mAb sequences of this clonal lineage, mAb10 and mAb22, were found in plasma at a percentage of 6.8% and 0.1% of all anti-MSP1 IgG detected, respectively, making it the fifth largest clonotype in the circulation. Despite having highly similar heavy chain variable regions, including eight shared amino acid substitutions and a long HCDR3 sequence (21 amino acid residues, **Supplementary figure 2**), mAb10 and mAb22 had different light chains. Like all other members of the clonal lineage (total n = 9), mAb22 had a lambda light chain, while mAb10 contained a kappa light chain. mAb10 was more abundant in plasma (6.8%), but was the only member of the MBC lineage with a kappa light chain. The opposite was observed for the lambda light chain variants, found at 0.1% of all anti-MSP1 plasma IgG, but representing eight out of nine members of the MSP1-specific MBC lineage. This difference in relative abundance of lineage members in different B cell compartments may be related to the preferential differentiation of high-affinity B cells to plasma cells and lower-affinity B cells to MBCs (33). For mAb10 and mAb22, it would therefore be expected that mAb10 has higher antigen-binding affinity. We determined binding affinity of mAb10 and mAb22 to MSP1 using a chaotropic ELISA with urea and observed that mAb10 indeed showed higher binding affinity to MSP1 than mAb22 (**Figure 6A**). The observation that mAb10 and mAb22 had different light chains suggests that antigen binding by these mAbs is dominated by their heavy chain. To test this, we expressed mAb22 with light chains from unrelated non-MSP1-binding antibodies and observed that it was still reactive with MSP1_3D7_, albeit with lower binding affinity (**Figure 6B**). These results suggest that the heavy chain of mAb22 (and presumably mAb10) is sufficient for binding to MSP1, but that its light chain is important for optimal binding to antigen. This raised the question whether the difference in binding affinity between mAb10 and mAb22 is caused by the different light chains used by the two mAbs. For each mAb, we therefore compared binding affinity between the antibody expressed with its own light chain and a chimeric antibody in which the light chain was swapped. Expression of mAb10 with the mAb22 light chain resulted in a reduction of binding affinity, while the binding affinity of mAb22 was unchanged when expressed with the mAb10 light chain (**Figure 6C**). These results suggest that the light chain of mAb10 may play a role in the increased binding affinity of mAb10 over mAb22, but is dependent on the mAb10 heavy chain for this effect. Interestingly, despite higher binding affinity, mAb10 showed reduced activity in a growth inhibition assay as compared to mAb22 (**Figure 2C**). Collectively, these results highlight characteristics of the individual members of an MSP1-specific clonal B cell lineage that may have influenced their selection and fate.

**Figure 6:**
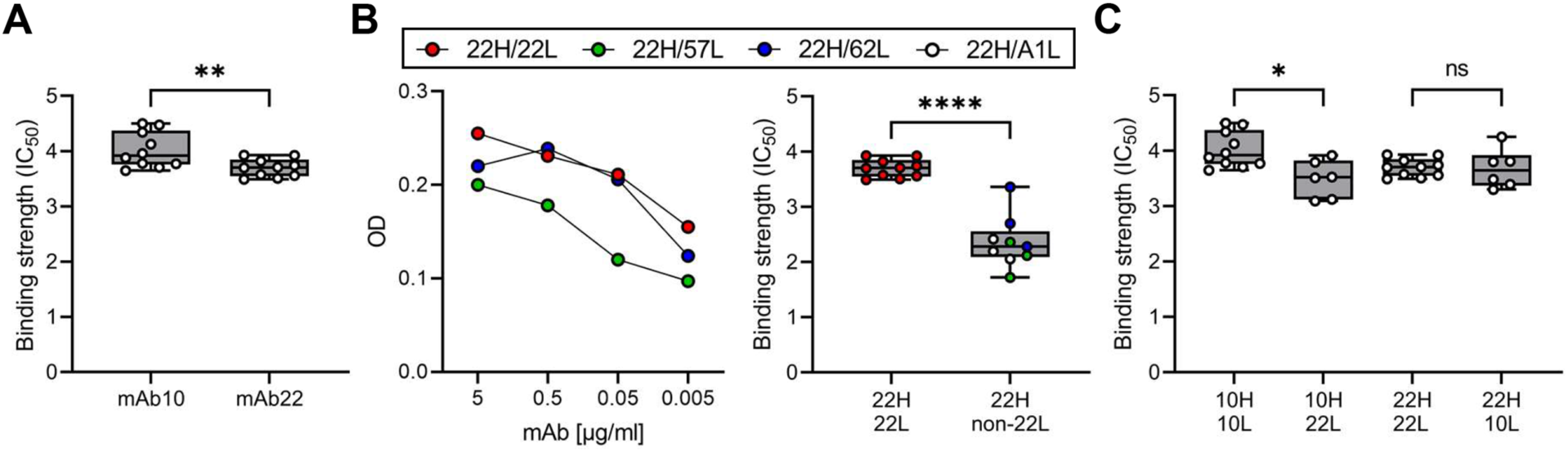
Binding affinity of mAb10 and mAb22 to MSP1_3D7_. **A)** The binding strength of mAb10 and mAb22 as determined by chaotropic ELISA. Binding strength (IC_50_) is defined as the concentration of urea that results in loss of 50% of mAb binding. A total of 10 replicates from three independent experiments are shown. Differences were tested for statistical significance using the Wilcoxon matched-pairs signed rank test, with pairs were defined as antibodies analyzed on the same plate. **B)** The binding of mAb22 and antibodies that consisted of the heavy chain of mAb22 and a light chain of an unrelated, non-MSP1-reactive antibody to MSP1_3D7_ by regular ELISA (left) and chaotropic ELISA (right). Differences in binding strength between replicates of mAb22 (n = 10) and chimeric antibodies (n = 9, three replicates for each of three unrelated light chains that were combined for analysis) were tested for statistical significance using the Mann Whitney test. **C)** The binding strength of mAb10 and mAb22 (n = 10 replicates each) expressed with their own light chains or chimeric mAbs in which the light chains were swapped (n = 6 replicates each). In all panels, data points are the average of three technical replicates. Differences between antibodies with the same heavy chain but a different light chain were tested for statistical significance using a Kruskal-Wallis test, followed by comparisons between select groups using Dunn’s post-hoc test, corrected for multiple comparisons. **** P < 0.0001; ** P < 0.01; * P < 0.05

## Discussion

The prevalence and magnitude of antibody responses against *P. falciparum* merozoite antigens have been extensively studied. However, much remains unknown about the molecular characteristics of anti-merozoite antibodies and the connections between the MBC and plasma cell compartments. This information will increase our understanding of how durable B cell immunity develops and how this may be harnessed for vaccine development. Here, we compared the phenotype and molecular characteristics of MBCs against the most abundant merozoite surface antigen, MSP1, between children and adults living in a region of high *P. falciparum* transmission in Uganda. In addition, we analyzed cross-strain reactivity among anti-MSP1 plasma IgM and IgG in both children and adults. Finally, we analyzed the overlap between the MBC and plasma antibody compartments in detail for one adult to better understand the processes that drive B cell selection and fate decisions.

Our results showed that children harbored a larger fraction of MSP1-specific IgM^+^ MBCs than adults, in line with a recent report (16). Interestingly, based on these observations, one might expect that the plasma anti-MSP1 antibody response would also be dominated by IgM, but this was not the case. In both adults and children, the level of anti-MSP1 in plasma was much lower than that of anti-MSP1 IgG. We recognize that it is difficult to directly compare IgM and IgG measurements due to potential differences between secondary antibodies used in these assays. However, our control samples from recovered COVID-19 patients demonstrate that we were able to measure IgM reactivity, yet we detected plasma IgM reactivity against MSP1 in only a small percentage of adults. These observations suggest that malaria-experienced children develop a strong IgG response against MSP1 that is not reflected in the MBC compartment. One explanation for the relative lack of IgG^+^ MBCs in children could be that the B cell response predominantly gives rise to short-lived IgG^+^ plasmablasts, whereas germinal center reactions are limited. Since germinal center responses are essential for the development of long-lived plasma cells and MBCs, this would explain the relatively quick waning of anti-parasite plasma IgG and MBCs in children in the absence of exposure, as seen in children living in malaria-endemic regions with a distinct dry season (18, 19). It has been shown that *P. falciparum* infections can lead to dysregulation of various components of the immune system that are essential for the formation of germinal centers, including dendritic cells and T follicular helper cells (34–37). In addition, the abundant development of plasmablasts during *Plasmodium* infection in rodents directly limited the generation of germinal centers as a result of nutrient depletion (38). In this scenario, IgM^+^ MBCs would mainly develop outside of germinal centers, which would be in agreement with the low levels of somatic hypermutation that we observed. An alternative explanation for our observation could be that anti-merozoite IgG^+^ MBCs are formed equally efficiently in both children and adults but undergo recall shortly after their formation in children as a result of repetitive *P. falciparum* infections with high parasitemia, whereas the lower antigenic load of asymptomatic infections in adults would allow for a longer lifespan of IgG^+^ MBCs. This theory would be in line with the increase in the percentage of merozoite-specific MBCs with age that is observed in individuals who live in malaria-endemic regions (19). In support of this theory is our observation that the few IgG^+^ MBCs isolated from children already had relatively high levels of amino acid substitutions, suggestive of multiple rounds of affinity maturation in germinal centers. It is also supported by our observation of a lack of clonal connections between IgG^+^ MBCs at two time points six months apart in two five-year old children living in a high transmission region in Uganda, while we did find clonally related sequences among IgM^+^ MBCs between the same two time points (Gonzales *et al.*, in press). These data could suggest that the IgG^+^ MBC compartment in children undergoes more rapid turnover than the IgM^+^ MBC compartment, or that the IgM^+^ MBC compartment has more self-renewing capacity (39). It is difficult to untangle various potential scenarios of how B cell responses develop and are influenced by the chronic and repetitive nature of *P. falciparum* infections. More longitudinal studies in the same individuals will be needed to track the fate of parasite-specific IgG^+^ MBCs after their formation.

We observed a low level of plasma IgM against both MSP1 and whole merozoites as compared to plasma IgG. This raises the question of how much IgM contributes to the control of *P. falciparum* infections. IgM responses against a variety of *P. falciparum* antigens have been reported in several studies, in particular in high transmission regions (40–43), but there is conflicting evidence on the durability of the IgM response and its role in parasite inhibition. Boyle *et al.* reported that IgM responses against MSP2 were sustained in children living in Kenya during a period with low parasite transmission (14). In contrast, Walker *et al.* concluded that IgM responses against five other merozoite antigens decreased rapidly following *P. falciparum* malaria in children living in Ghana, whereas IgG responses against these five antigens were maintained for a longer period of time (43). These conflicting observations may be the result of differences between the assays used to assess these antibody responses, or differences in the development of IgM responses under different transmission intensities or against different parasite antigens. An association between IgM reactivity against *P. falciparum* antigens, including MSP1_19_, AMA1, MSP3, and GLURP, and protection against malaria was reported by some (14, 44, 45), but not by others (46). When tested at the same concentration, purified plasma IgM from malaria-experienced individuals had equal opsonic phagocytosis activity and two-fold lower fixation capacity of C1q, the primary component of the classical complement pathway, as compared to plasma IgG, but induced nine-fold higher deposition of components of the membrane attack complex (14, 16). While these results suggest that IgM can indeed contribute to parasite inhibition, the concentration of IgM in plasma is almost one order of magnitude lower than that of IgG (average, 1.5 g/l for IgM versus 11 g/l for IgG, (47, 48)). Therefore, the relative contributions of IgM and IgG to parasite inhibition remain to be determined. Finally, a passive immunization study showed that IgG-depleted plasma from malaria-experienced adults had no effect on parasitemia in children with *P. falciparum* malaria, whereas treatment with IgG from the same individuals resulted in a dramatic decrease of parasite counts (49), suggesting that IgG is the main effector antibody responsible for parasite control.

The acquisition of high levels of amino acid substitutions in IgG^+^ MBCs from both children and adults suggests that these cells are the product of multiple rounds of affinity selection. We also detected several B cells with long HCDR3s, similar to broadly neutralizing antibodies against HIV (50), although this was not a universal characteristic of anti-MSP1 mAbs. In addition, we observed that children and adults had equally high IgG reactivity against MSP1 3D7, but that reactivity against MSP1 HB3 was lower in children. This suggests that the IgG response continues to broaden with subsequent *P. falciparum* infections, resulting in a larger fraction of IgG directed against conserved epitopes. The broadening of antibody responses, high mutational load, and expansion of clonal MSP1-specific B cell lineages is suggestive of strong selection pressure on the B cell response as a result of *P. falciparum* infections. We believe that our observations regarding mAb10 and mAb22, two members of an MSP1-specific IgG^+^ MBC lineage that share the same heavy chain but have different light chains, are additional evidence of strong forces on B cell selection and development. The difference in light chain usage between mAb10 and mAb22 raises the question how these mAbs are developmentally related. One possibility is that mAb10 and mAb22 were derived from a B cell precursor that underwent cell division upon heavy chain rearrangement in the bone marrow, giving rise to multiple daughter cells that each underwent light chain rearrangement independently. In this scenario, the shared amino acid mutations would be the result of convergent evolution selecting for higher affinity antigen binding. In a second scenario, a precursor B cell activated by antigen could have undergone secondary light chain rearrangement in the germinal center. In this case, the kappa light chain-bearing mAb10 would likely be evolutionarily closer to this precursor, and mAb22 (and all other MBCs in this lineage) would be derived from this secondary rearrangement event and have undergone further diversifying selection. We were unable to determine whether mAb10 and mAb22 developed as a result of convergent evolution or clonal lineage diversification. However, our results suggest that the light chains of these antibodies may affect antibody binding affinity. This could, on its own, have resulted in the preferential differentiation of the higher affinity mAb10 variant into a plasma cell and the lower affinity mAb22 variant into an MBC. Further studies into the relationship between the plasma cell and MBC compartments for MSP1 and other merozoite antigens are currently ongoing in our laboratories.

In conclusion, we performed an analysis of the MBC compartment and antibody responses against MSP1 in children and adults living under high *P. falciparum* transmission conditions. We observed that children predominantly harbor MSP1-specific IgM^+^ MBCs, while the MBC response had shifted to IgG^+^ B cells in adults. In contrast to this difference in isotype among MSP1-specific MBCs, both children and adults demonstrated strong anti-MSP1 plasma IgG responses, while anti-MSP1 plasma IgM responses were minimal. IgG^+^ MBCs carried high levels of somatic hypermutations and clonal expansion, suggestive of ongoing B cell selection over the course of sequential *P. falciparum* infections. Finally, we directly compared the overlap between the MSP1-specific MBC compartment and anti-MSP1 plasma IgG, revealing that the same molecular characteristics observed in the MBC compartment were also dominant in the plasma IgG compartment. Collectively, our results provide new insights into the development of B cell responses against *P. falciparum*, in particular about the similarities and differences between MBC and antibody responses.

## Materials and methods

### Ethics approval statement

Participants were enrolled in the Tororo Child Cohort or the Program for Resistance, Immunology, Surveillance, and Modeling of Malaria (PRISM) Cohort, and provided written consent for the use of their samples for research. These cohort studies were approved by the Makerere University School of Medicine Research and Ethics Committee (SOMREC) and the University of California, San Francisco Human Research Protection Program & IRB. The use of cohort samples was approved by the Institutional Review Board of the University of Texas Health Science Center at San Antonio. Donors 1 and 2 were anonymous blood donors at Mbale regional blood bank in Eastern Uganda, who consented to the use of their blood for research. The use of samples from anonymous blood donors was considered not human research by the Institutional Review Board of the University of Texas Health Science Center at San Antonio.

COVID-19 samples used in this study were received de-identified from the University of Texas Health San Antonio COVID-19 Repository. This repository was reviewed and approved by the University of Texas Health Science Center at San Antonio Institutional Review Board. All study participants provided written informed consent prior to specimen collection for the repository to include collection of associated clinical information and use of left-over clinical specimens for research. The COVID-19 Repository utilizes an honest broker system to maintain participant confidentiality and release of de-identified data or specimens to recipient investigators.

### Participants

Malaria-experienced participants were residents of Eastern Uganda, a region with extremely high malaria transmission intensity (annual entomological inoculation rate in the region estimated at 125 infectious bites per person per year (51). In this region, children between five and ten years of age start to develop protective immune responses against malaria but are typically still susceptible to disease. Children above ten years of age and adults have developed protective immune responses, evidenced by no signs of clinical malaria despite high exposure, demonstrated by household entomological and epidemiological measures, including documented asymptomatic parasitemia.

### MSP1 expression and tetramer synthesis

To produce C-terminal biotinylated 3D7 MSP1, human Expi293F cells (Thermo #A14635) were cultured, passaged, and transfected with the plasmids MSP1-bio (Addgene #47709) and secretedBirA-8his (Addgene #32408) at a 4:1 (w/w) ratio according to Thermo’s protocol. Both plasmids were a kind gift from Gavin Wright (52, 53). Biotin (Thermo #PI21336) was added to a final concentration of 100 µM immediately after adding the transfection mix. Cell were cultured in either non-baffled polycarbonate flasks with a vent cap (Fisher #PBV12-5) or in glass Erlenmeyer flasks loosely covered with aluminum foil to allow for gas exchange. Cells were successfully passaged in volumes as low as 5 ml. Absence of mycoplasma contamination was confirmed using the MycoAlerta Plus mycoplasma detection kit (Lonza #LT07705). Culture supernatants were collected 5 – 7 days post-transfection by centrifuging the culture at 4,000 × g for 25 min. at RT. A 10 kDa cutoff Protein Concentrator PES (Thermo #88527) was used (5,000 × g at 4⁰C) to exchange culture medium containing free biotin for PBS (pH 7.2) (> 100,000 dilution) and to concentrate the protein to a final volume of 0.5 – 1 ml. The 3D7 MSP1 protein was mixed with 6 – 12 volumes of PBS (pH 5.5) in a final volume of six ml and was subsequently loaded onto gravity flow columns (Thermo #29924) containing CaptAvidin agarose (Thermo #C21386) for purification. After three washes with PBS (pH 5.5) and five 6 ml elutions with PBS (pH 10.5), the elutions were pooled (30 ml) and the pH was immediately neutralized by adding 12 ml PBS (pH 5.5). After concentrating, the protein was quantified using the Coomassie Plus (Bradford) Assay Kit (Thermo #23236) on a NanoDrop One spectrophotometer, according to the manufacturer’s instructions, visualized by SDS-PAGE (**Supplementary Figure 3A**), diluted to 1 mg/ml, aliquoted, and stored at -70°C.

Since each streptavidin molecule has the ability to bind four biotinylated MSP1 molecules, MSP1 tetramers were made by incubating MSP1 in a tube revolver (Thermo #88881001) at 40 rpm and RT for 30 min. with streptavidin-PE (Thermo #S866) at a 6:1 molar ratio. After this incubation, the tetramers were washed with PBS (pH 7.2) using a Vivaspin centrifugal concentrator (Sartorius #VS0141) three times for 5 min. at 15,000 × g at RT. To make decoy tetramers, streptavidin-PE was first conjugated to Alexa-fluor 647 (Thermo #A20186) per manufacturer’s instructions. This double-conjugated streptavidin was then coupled to *R. norvegicus* CD200 (Addgene #36152 (53)) as described above.

### SDS-PAGE

As a quality control step, purified MSP1 and antibodies were visualized on a polyacrylamide gel. Purified samples were mixed with Laemmli buffer and NuPage sample reducing agent (Thermo #NP0004; not added for monoclonal antibodies), incubated at 85°C for two min. The samples were run on a 4 – 12% Bis-Tris gel (Thermo #NP0321BOX) with MOPS running buffer (Thermo #NP0001) at 200 V for 50 min. The proteins were stained using Imperial Coomassie protein stain (Thermo #PI24615) per manufacturer’s instructions.

### MSP1-specific B cells isolations

Cryopreserved peripheral blood mononuclear cells (PBMCs) from malaria-experienced children and adults were thawed in a water bath at 37°C and immediately mixed with pre-warmed thawing medium (IMDM Glutamax (Thermo #31980030) supplemented with 10% heat-inactivated fetal bovine serum (FBS) of US origin (Sigma #TMS-013-B) and 33 U/ml universal nuclease (Thermo #88700)) and then centrifuged (5 min. at 250 × g and RT). The cell pellet was resuspended in thawing medium and viable cells were counted by adding 10 µl filtered 0.2% trypan blue in PBS to 10 µl of the cell suspension on a Cellometer Mini (Nexcelom) automated cell counter. Next, cells were pelleted by centrifugation (5 min. at 250 × g and RT) and resuspended in isolation buffer (PBS supplemented with 2% heat-inactivated FBS and 1 mM EDTA) at 50 million live cells/ml and filtered through a 35 µm sterile filter cap (Corning #352235) to break apart any aggregated cells. B cells were isolated using StemCell’s EasySep Human B Cell Isolation Kit (#17954) according to manufacturer’s instruction. After washing with PBS, the isolated B cells were incubated with 1 µl LIVE/DEAD Fixable Aqua Dead Cell Stain Kit (Thermo #L34965) per 1 ml cell suspension, per manufacturer’s instructions. After washing the B cells with cold PBS and resuspending them in 50 µl cold PBS with 1% bovine serum albumin (BSA) (Sigma #A7979), the cells were first stained with 40 nM of decoy tetramer (10 min. in the dark on ice) and then with 20 nM of MSP1 tetramer (30 min. in the dark on ice), followed by a wash with 1 ml of cold PBS/1% BSA (5 min. at 250 × g and RT). Tetramer-bound B cells were selected using StemCell’s EasySep Human PE Positive Selection Kit (#17664) and subsequently stained on ice for 30 min. with an antibody panel against B cell surface markers (**Supplementary table 4**).

UltraComp eBeads (Thermo #01222242) were used to prepare compensation controls for each fluorophore per manufacturer’s instructions. Before acquisition on a BD FACSAria II cell sorter, the cells were washed with 3 ml of cold PBS with 1% BSA (5 min. at 250 × g and 4°C), diluted to 20 – 30 million cells/ml in PBS with 1% BSA, and filtered into a FACS tube with filter cap. Lymphocytes were gated using forward and sideward scatter, followed by doublet exclusion and gating on live cells. MSP1-specific mature IgG^+^ and IgM^+^ B cells (CD19^+^, CD20^+^) were gated (PE^+^, AF647^-^) and single cells were sorted into 100 µl IMDM/Glutamax/10% FBS in a well of a 96-well plate (Corning #353072). One day prior to the sort, each well was seeded with 30,000 adherent, CD40L-expressing 3T3 cells (kind gift from Dr. Mark Connors, NIH) in 100 µl IMDM/Glutamax/10% FBS containing 2× MycoZap Plus-PR (Lonza #VZA-2021), 100 ng/ml human IL-2 (GoldBio #1110-02-50), 100 ng/ml human IL-21 (GoldBio #1110-21-10), 5 µg/ml TLR9-activator ODN2006 (IDT DNA, sequence TCGTCGTTTTGTCGTTTTGTCGTT), and 60 µg/ml transferrin (Sigma #616424) to promote expansion and differentiation of B cells into antibody-secreting cells (54, 55). After incubation at 37°C and 8% CO_2_ for two weeks, the wells were screened for the production of IgM or IgG by enzyme-linked immunosorbent assay (ELISA).

### Enzyme-linked immunosorbent assays

To detect IgG and IgM, 96-well ELISA plates (Corning #3361) were coated with either goat anti-human IgG (Sigma #I2136) or IgM (Sigma #I1636) antibody at a concentration of 4 and 8 µg/ml, respectively, diluted in PBS, at a total volume of 100 µl per well. After a one hour incubation at 37°C or O/N at 4°C, each well was washed once using slowly running (approximately 900 ml / min.) deionized water. This washing method resulted in significantly higher specificity than other methods tested in the lab (using a plate washer with water or PBS containing 0.1% tween-20, or a squeeze bottle filled with PBS containing 0.1% tween-20). All subsequent washes were performed this way. 150 µl blocking buffer (one-third Non-Animal Protein (NAP)-Blocker (G-Biosciences #786-190P) and two-thirds PBS) was added to each well to prevent non-specific binding. After one hour of incubation at 37°C, the wells were washed three times and 50 µl B cell culture supernatant diluted 1:1 in dilution buffer (1% NAP Blocker in PBS; total volume 100 µl) was added per well. Plates were incubated for two hours at 37°C and washed five times. Then, either 100 µl 1:2500 diluted (1% NAP Blocker in PBS) HRP-conjugated anti-human IgG antibody (BioLegend #410902) or 1:5000 HRP-conjugated anti-human IgM antibody (Sigma #AP114P) was added to each well. After incubation for one hour at 37°C and three washes, HRP activity was detected using 50 µl TMB (Thermo #PI34024). Plates were incubated in the dark at RT and the oxidation reaction was stopped by adding 50 µl 0.18M H_2_SO_4_ (Fisher #FLA300-212) per well when the negative controls (wells that received buffer when test wells received culture supernatant) started to color. Absorbance was measured at 450 nm using a BioTek Synergy H4 microplate reader. A human IgG (Sigma #I2511) or IgM (Sigma #I8260-1MG) standard curve (ten three-fold serial dilutions starting at 20 µg/ml) was used to quantify samples. Wells with values >27 ng/ml were considered positive. This cutoff was determined based on our observation that the amplification of heavy and light chain variable regions always failed from cultures with a lower concentration.

ELISAs to confirm reactivity of MSP1-specific antibodies were performed as described above with the following modifications. Plates were coated with 50 µl in-house produced MSP1_3D7_ per well at a concentration of 16 µg/ml (0.8 µg/well). Coated plates were incubated for 1 hour at 37⁰C or overnight at 4°C and all subsequent incubations were done at RT instead of 37°C. To prevent non-specific binding, the wells were blocked with 200 µl PBS containing 0.1% tween-20 and 3% non-fat milk powder (SACO), which significantly increased specificity of the assay (compared to NAP blocker). After discarding the blocking buffer from the wells, the plates were not washed. Purified antibodies were tested at a final concentration of 2.5 µg/ml in 100 – 200 µl in PBS containing 0.1% tween-20 and 1% non-fat milk powder. The plates were washed six times prior to adding the detection antibody, and four times prior to adding TMB substrate. To analyze binding affinity of monoclonal antibodies to MSP1_3D7_, chaotropic ELISAs were performed similar to the above descriptions, with the following modifications. Wells were coated with 0.2 µg protein and the final concentration of the antibodies that were tested was 0.5 µg/ml. Following the incubation with anti-MSP1 antibodies, the plates were washed four times. Then, urea (Fisher #U15-500) was added to wells at the following concentrations: 0, 1, 2, 3, 4, 5 and 8M (in 100 µl PBS with 0.1% tween-20 and 1% milk powder). After a 15-minute incubation at RT, plates were washed four times. The IC_50_ (the molar concentration of urea required to reduce antibody binding to MSP1 by 50%) was calculated using non-linear regression analysis in GraphPad Prism 9. Urea concentrations were log-transformed prior to analysis. OD values for each technical replicate were normalized by setting the smallest OD to 0% and the largest OD to 100%. The IC_50_ values of three technical replicates were averaged to obtain the final IC_50_ for an experiment.

ELISAs to detect antibody reactivity against merozoites were performed as described for MSP1-specific antibodies above with the following modifications. Free merozoites were coated at 500,000 merozoites per well in 100 µl PBS, followed by overnight incubation at 4°C. Merozoite count was estimated based on culture parasitemia. Plasma samples were tested at a 1:200 dilution in a total volume of 100 µl.

### Amplification of antibody heavy and light chain variable regions

MSP1-specific B cells that successfully expanded in culture were collected by centrifugation (5 min. at 250 × g and RT) and stored at -70°C in 50 µl Tri-Reagent (Zymo #R2050-1-200). Heavy and light chain variable regions were amplified from MSP1-specific B cells by cDNA synthesis and a series of PCR reactions, shown in **Supplementary figure 4A**. All primer sequences can be found in **Supplementary table 5**. mRNA was isolated using Zymo’s Direct-zol RNA Microprep kit (#R2060), eluted in 15 µl elution buffer and then mixed with 0.7 µl reverse primer (10 µM, 200 mM final concentration (f/c) in 35 µl PCR reaction volume) specific for the IgG, or IgM, heavy chain (primers #7 and #297) plus 0.7 µl light chain specific reverse primers (10 µM): #108 and #109,), and incubated for two minutes at 65°C. Single stranded cDNA was synthesized immediately by adding 0.7 µl SMARTScribe reverse transcriptase (100 U/µl, f/c 2 U/µl, Takara Bio #639537), 7 µl First-Strand buffer (f/c 1×), 7 µl DTT (20 mM, f/c 4 mM), 0.7 µl dNTPs (10 mM each, f/c 200 µM each, Sigma #DNTP-10), 1.75 µl RNase OUT (40 U/µl, f/c 2 U/µl, Thermo #10777019), 0.7 µl template switch oligo (TSO) (10 µM, f/c 200 nM, IDT DNA; #110, **Supplementary table 5**), nuclease-free water till 35 µl and subsequent incubation at 42°C for 2 hours. The TSO was designed with two isodeoxynucleotides at the 5’ end to prevent TSO concatemerization and three riboguanosines at the 3’ end for increased binding affinity to the appended deoxycytidines (property of the Takara reverse transcriptase) (56, 57). The single-stranded cDNA was immediately purified using Zymo’s RNA Clean & Concentrator kit (#R1016) using Zymo’s appended protocol to purify fragments >200 nucleotides and was eluted in 10 µl elution buffer. This critical clean-up step ensured that any unused TSO was removed, preventing it from inhibiting the subsequent PCR reactions by serving as template for the forward primer. Immediately after, heavy and light chain variable regions were amplified by PCR in one reaction mix using 8.5 µl purified cDNA, 10 µl AccuStart II PCR SuperMix (QuantaBio #95137), 0.9 µl 10 µM forward primer #106 (f/c 0.45 µM, **Supplementary table 5**), and 0.2 µl of the reverse primers (10 µM) used to synthesize the cDNA (#7, #297, #108, and #109, each at f/c 0.1 µM). Cycling conditions were 94°C for 3 min., 35 cycles of 30 sec. at 94°C, 30 sec. at 55°C and 35 sec. at 72°C, followed by 5 min. at 72°C. A second, nested amplification was required to obtain enough amplicon DNA, and was done separately for heavy chain, kappa light chain, and lambda light chain variable regions, using AccuStart II PCR SuperMix, and 2 µl of the first, unpurified PCR as template in a total reaction volume of 20 µl. Mixes of primers (**Supplementary figure 4A**) as described by Hua-Xin Liao *et al*. (58) were used for this second PCR, with a final concentration of 0.1 µM for each individual primer. Reverse primer #67 was added for the heavy chain variable region PCR to allow for amplification of variable regions originating from IgG_2_, IgG_3_ and IgG_4_ mRNA, in addition to #30 which was specific for IgG_1_. Cycling conditions were as described above, except for the extension step (shortened to 30 sec.) and the annealing step, which was 30 sec. at 60°C for the IgG1 heavy chain variable region, 30 sec. at 63⁰C for the IgM heavy chain variable region, and 30 sec. at 50°C for the light chain variable regions.

Linear IgG expression cassettes (58) were synthesized by PCR using 3 overlapping DNA fragments: a promoter (705 bp), a variable region and a constant region (IgG_1_ heavy chain: 1188 bp; pentameric IgG_1_ heavy chain: 1326 bp; kappa light chain: 568 bp; lambda light chain: 534 bp). Details about the assembly of linear expression cassettes are described by Liao *et*. *al*. (58). All fragments were amplified in a 100 µl reaction using 20 ng plasmid template (HV0023 – HV0026 (58) or the IgG_1_ expression plasmid containing the IgM multimerization sequence (see below)), 4 µl forward primer (10 µM), 4 µl reverse primer (10 µM), and QuantaBio AccuStart II PCR supermix (2×) using the following cycling program: 94⁰C for 3 min., 35 cycles of (94⁰C for 30 sec., 68⁰C for 30 sec., and 72⁰C for 40 – 75 sec. (1 min. per 1000 bp)) and 72⁰C for 5 min. Primers 53 and 54 were used for the promoter fragment, 55 and 58 for the heavy chain constant region fragment, 56 and 58 for the kappa light chain constant region fragment, 57 and 58 for the lambda light chain constant region fragment, and 55 and 494 for the pentameric IgG_1_ heavy chain constant region fragment (**Supplementary table 5**). The overlapping PCRs were done as follows. The cycling program was the same for all overlapping PCRs: 98⁰C for 1 min., 30× (98⁰C for 20 sec., 68⁰C for 15 sec., 72⁰C for 60 sec.), 72⁰C for 10 min. Two ng of each fragment (promoter, constant region, variable region) was used as template in a 25 µl PCR reaction with 2× KAPA HiFi Hot Start Ready Mix (Roche #KK2602) and 1 µl (10 µM) of the following forward and reverse primers: 50 and 51 for the IgG1 heavy chain, 50 and 52 for the kappa and lambda light chain, and 50 and 469 for the IgG_1_ heavy chain with IgM multimerization domain (**Supplementary table 5**). The final size of the linear expression cassettes was ∼2300 bp for the IgG1 heavy chain, ∼2400 bp for the IgG1 heavy chain with IgM multimerization domain, and 1600 bp for the kappa and lambda light chains (**Supplementary figure 4B**). All linear expression cassettes were purified and sequence verified by Sanger sequencing. Variable region sequences were analyzed with IMGT/V-QUEST (28) using default settings to identify V(D)J gene usage and amino acid substitutions.

### Generation of antibody expression plasmids

Antibody variable regions were cloned into expression plasmids from Invivogen (#pfusess-hchg1, #pfuse2ss-hclk, #pfuse2ss-hcll2). The variable heavy and light chain regions were amplified from the linear expression cassettes (2 µl at 1 ng/µl) using 10 µl NEB Q5 Hot Start HiFi PCR master mix (#M0494S), 6 µl nuclease-free water and 1 µl sequence-specific F and R primer (10 µM, f/c 500 nM) that were based on the results of analysis using IMGT/VQUEST (28). These primers introduced restriction sites (EcoRI & NheI for hchg1, EcoRI & BsiWII for hclk, and EcoRI & AvrII for hcll2). Annealing temperatures were primer sequence dependent and were calculated using NEB’s Tm calculator to match the salt concentration in their buffer. In an attempt to express the variable regions from IgM^+^ B cells as a multimer (pentamer/hexamer mix) instead of a monomer, we modified the IgG_1_ heavy chain expression plasmid (15). The IgM multimerization sequence PTLYNVSLVMSDTAGTCY (CCAACGCTCTATAATGTCTCTTTGGTTATGTCCGACACAGCCGGTACCTGCTAT) was cloned into the IgG_1_ expression vector at the C-terminus of the open reading frame, immediately in front of the stop codon, and the leucine at position -139 relative to the proline in the multimerization sequence was changed into a cysteine. Every plasmid was Sanger sequence-verified prior to using it as expression vector.

### Antibody expression and purification

For small scale screening, one ml Expi293F cell cultures in a 6 well plate were transfected with heavy and light chain linear expression cassettes (1:1 molar ratio) according to the manufacturer’s instructions for 25 – 30 ml cultures (also at 125 rpm). Heavy and light chain antibody expression plasmids were used at a molar ratio of 1:2 to transfect 5 ml cultures. The antibodies were purified from the culture supernatant 4 – 6 days later using protein G magnetic beads (Promega #G7472). Purified antibodies and antibody elution buffer (5 parts elution buffer (100 µM glycine-HCl, pH 2.7) and 1 part neutralization buffer (2M Tris buffer, pH 7.5)) were buffer exchanged to PBS using 100 kDa cutoff Protein Concentrators (Thermo #88523). The samples were diluted > 50,000 × in PBS by repeated centrifugation at 4,000 × g and 4⁰C. Purified antibodies were quantified using the Coomassie Plus (Bradford) Assay Kit (Thermo #23236) on a NanoDrop One spectrophotometer, according to the manufacturer’s instructions, and visualized on SDS-PAGE gel with a standard amount of BSA to confirm protein size and purity (**Supplementary figure 3B**).

### Parasite culture and merozoite isolation

*P. falciparum* 3D7 strain parasites were cultured (59) in human AB^+^ erythrocytes (Interstate Blood Bank, Memphis, TN, USA) at 3 – 10% parasitemia in complete culture medium (5% hematocrit). Complete culture medium consisted of RPMI 1640 medium (Gibco #32404014) supplemented with gentamicin (45 µg/ml final concentration; Gibco #15710064), HEPES (40 mM; Fisher #BP3101), NaHCO_3_ (1.9 mg/ml; Sigma #SX03201), NaOH (2.7 mM; Fisher #SS266-1), hypoxanthine (17 µg/ml; Alfa Aesar #A11481-06), L-glutamine (2.1 mM; Corning #25005Cl), D-glucose (2.1 mg/ml; Fisher #D16-1), and 10% heat-inactivated human AB^+^ serum (Valley Biomedical #HP1022). Parasites were cultured at 37°C in an atmosphere of 5% O_2_, 5% CO_2_, and 90% N_2_. Before use in cultures, 12.5 ml packed erythrocytes were washed twice with 10 ml cold incomplete medium (complete culture medium without human serum) and pelleted between each wash by centrifugation at 500 × g for 8 min. at 4°C (max. acceleration and weakest break). Washed erythrocytes were resuspended in 2 volumes of complete medium and stored at 4°C.

Parasites were synchronized to the ring stage by treatment with 5% D-sorbitol (60) (Fisher #S459-500). Cultures containing high percentages of ring-stage parasites were centrifuged at 250 × g for 5 min. at RT. Pelleted erythrocytes were resuspended in 10 volumes of 5% D-sorbitol in MQ water, vortexed for 30 sec. and incubated for 8 min. at 37°C. The cells were then washed with 5 volumes of complete culture medium (250 × *g* for 5 min. at RT) and resuspended in complete culture medium at 5% hematocrit and cultured as described above. To obtain tightly synchronized parasites, sorbitol treatments were performed twice, 14 hours apart.

Infected erythrocytes containing parasites in the late-trophozoite and schizont stages were isolated from culture by magnetic separation (61, 62). Late-stage parasites were separated from uninfected and ring-infected erythrocytes with a SuperMACS II Separator (Miltenyi #130-044-104). The magnet was assembled with a D column (Miltenyi #130-041-201) according to manufacturer’s instructions. The column was equilibrated with 200 ml incomplete medium. An additional 50 ml incomplete medium was added to the column through the side syringe to remove air bubbles possibly remaining in the column matrix. A 22 G needle (BD #305155) was attached to the stopcock to serve as a flow restrictor. For safety purposes the plastic protective sheath remained on the needle after cutting the end to allow flow of the liquid without exposing the tip of the needle. Approximately 100 – 200 ml of synchronized parasite culture (5 – 10% parasitemia, 5% hematocrit) 24 – 27 hours following the second sorbitol treatment (majority of parasites in the early segmented schizont stage, 4 – 6 nuclei visible by Giemsa staining) were used for merozoite isolation. After passing the parasite culture through the column, the column was washed from the top with incomplete medium until the flowthrough was clear (usuall ∼100 ml). Next, the column was washed with a total of 150 ml incomplete medium (50 ml from the side and 100 ml from the top). Erythrocytes containing late-stage parasites with high paramagnetic hemozoin levels are preferentially retained in the column matrix while attached to the magnet (63) allowing for separation of late-stage parasites from uninfected erythrocytes and early-stage parasites. The column was removed from the magnet and 60 ml incomplete medium was used to elute the erythrocytes from the column matrix. The erythrocytes were pelleted by centrifugation at 250 × *g* for 5 min. at RT and were resuspended in 3 ml complete culture medium. Infected erythrocytes were incubated with E64 (10 µM final concentration, Sigma #324890-1MG) for 8 hours at normal culture conditions to allow the parasites to develop into fully segmented schizonts while preventing egress from the erythrocytes. Infected erythrocytes containing schizonts were then pelleted by centrifugation at 1,900 × *g* for 8 min. at RT and the supernatant containing E64 was removed. A thin smear from the pellet was Giemsa stained and merozoite yield was assessed by counting the number of fully segmented schizonts present. The pellet was resuspended in 4 ml incomplete medium. Merozoites were released from the erythrocytes by passing them through a 1.2 µm syringe filter (Pall #4190) and were subsequently pelleted by centrifugation at 4,000 × *g* for 10 min. at RT. On average, 5 × 10^7^ merozoites were collected per 25 ml of synchronized culture. The merozoites were resuspended in PBS and stored at 4°C for up to one day until used for the ELISA.

### Immunofluorescence assay

All steps of the immunofluorescence assay were done at RT. A thin blood smear was made on a microscopy slide from a 1 µl drop of E64-treated schizont culture. After drying for 30 sec., the cells were fixed by loading 1 ml of 4% paraformaldehyde (Electron Microscopy #15710) on the slide and incubating it for 30 min. The fixed cells were then washed three times with 1 ml PBS. Following the washes, the cells were permeabilized with 0.1% Triton-X (Fisher # BP151) in PBS and incubated for 30 min. The cells were then washed for an additional three washes using PBS. The slide was treated with blocking buffer (2% BSA, 0.05% Tween-20, 100 mM Glycine, 3 mM EDTA and 150 mM NaCl in PBS) for one hour. MSP1-specific mAbs were added to the cells at 1 µg/ml in 500 µl blocking buffer and incubated for one hour. Samples were then washed again three times using PBS. Goat anti-human IgG conjugated to FITC (Thermo #A18830) secondary antibody was diluted 1:1000 (1.5 µg/ml) in blocking buffer and then added to the smear to incubate for one hour in the dark. Samples were again washed three times using PBS in the dark and then allowed to air dry for one hour in the dark. Slides were mounted using 10 µl ProLong Glass mounting medium containing NucBlue Stain (Thermo #P36985) and sealed with a cover slip. Samples were imaged using a Zeiss Axio Imager Z1 with Zen Blue software.

### Luminex assay

100 pmol MSP1_3D7_, MSP1_Dd2_, MSP1_HB3_, and SARS-CoV-2 receptor binding domain were coupled per 1 × 10^6^ MagPlex microspheres (Luminex, #MC10025-ID) using the Luminex protein coupling kit (#40-50016) per manufacturer’s instructions. All subsequent steps were done at RT and the beads were protected from light using aluminum foil. Coupled beads were pooled, resuspended in buffer A (PBS with 0.05% Tween 20 (Fisher #BP337), 0.5% BSA (Sigma #A7979), 0.02% sodium azide) and plated at 1000 beads per well for each protein in a black, flat-bottom 96 well plate (Bio-Rad #171025001). The beads were washed once. All washes were done with 100 µl PBST (PBS with 0.05% Tween 20) using a handheld magnetic washer (Bio-Rad #171020100). The incubation time on the magnet was always 2 min. Next, the beads were incubated with 50 µl purified anti-MSP1 antibody (diluted to 1 µg/ml using buffer B) or B cell culture supernatant (diluted 1:1 with buffer B) for 30 min. with constant agitation (500 rpm, 2.5 mm orbital diameter). Buffer B (0.05% Tween-20, 0.5% BSA, 0.02% sodium azide, 0.1% casein (Sigma #C7078), 0.5% PVA (Sigma #P8136) and 0.5% PVP (Sigma #PVP360), 15 µg/ml *E. coli* lysate) was prepared a day prior to the assay since it required the chemicals to dissolve O/N. On the day of the assay, *E. coli* lysate (MCLAB #ECCL-100) was resuspended in MQ water and added to a final concentration of 15 µg/ml. Prior to use, buffer B was centrifuged at 10,000 × g for 10 min. After three washes, 50 µl secondary antibody diluted in buffer A (PE anti-human IgG (1:200 dilution; Jackson ImmunoResearch #109-116-098)) or PE anti-human IgM (1:80 dilution; BioLegend #314507)) was added per well. After 30 min. incubation with constant agitation, the beads were washed three times and subsequently incubated in 50 µl buffer A for 30 min. with constant agitation. After one final wash, the beads were resuspended in 100 µl PBS and fluorescence intensity was measured using a calibrated and validated Bio-Rad Bio-Plex 200 machine.

### Growth inhibition assay

*P. falciparum* isolate 3D7 parasites were pre-synchronized at the ring stage with a 5% D-sorbitol (Fisher #S459-500) treatment as described above, followed four days later by two additional 5% D-sorbitol treatments 14 hours apart (60). At the late trophozoite / early schizont stage (24 hours after the third D-sorbitol treatment), parasitemia was determined by inspection of a Giemsa-stained blood smear. The smear was also used to confirm correct parasite staging. Immediately after, 20 µl of each antibody (1 mg/ml in PBS) was added to wells containing 30 µl complete medium in a black clear bottom 96-well plate (Corning #3603). A monoclonal antibody specific for apical membrane antigen 1 (AMA1) was used as a positive control (BEI #MRA-481A). Antibody elution buffer (100 mM glycine-HCl, pH 2.7) that was buffer exchanged to PBS alongside purified antibodies (see “Antibody expression and purification” above) was used as a negative control. Fifty µl parasite culture (1% parasitemia and 2% hematocrit) was then added to wells containing antibody or negative control. Uninfected erythrocytes (2% hematocrit) were used to determine the background signal. The plate was then incubated at standard parasite culture conditions (described above) for 48 hours before being transferred to a -70°C freezer. After overnight incubation of the plate at -70°C, SYBR green dye (Invitrogen #S7585) was added to lysis buffer (20 mM Tris-HCl (pH7.5), 5 mM EDTA, 0.008% saponin (Sigma # 558255100GM), 0.08% Triton X-100 in MQ water) at 0.2 µl dye per ml of lysis buffer. One hundred µl SYBR green lysis buffer was added to each well and the plate was incubated in the dark at 37°C for 3 – 6 hours.

Fluorescence (excitation = 495 nm, emission = 525 nm, cutoff = 515 nm) was measured with a BioTek Synergy H4 plate reader. The instrument was programmed to read the plate from the bottom after mixing for 5 sec. The average background fluorescence value was subtracted from the fluorescence signal of the wells with infected cells. Percent growth inhibition was expressed as the reduction in fluorescence signal in wells incubated with antibody as compared to the negative control.

### Plasma antibody proteomics

Total IgG was isolated from 1 ml plasma using Protein G Plus Agarose (Thermo #22851) affinity chromatography and cleaved into F(ab’)_2_ fragments using IdeS protease. MSP1-specific F(ab’)_2_ was isolated by affinity chromatography using 1 mg recombinant MSP1_19_ produced in *E. coli* (Meridian Life Sciences, #R01603 and #R01604) coupled to 0.05 mg dry NHS-activated agarose resin (Thermo #26196) as follows. F(ab’)_2_ (10 mg/ml in PBS) was rotated at 8 rpm with antigen-conjugated affinity resin for 1 hour at RT, loaded into 0.5 ml spin columns (Thermo #89868), washed 12× with 0.4 ml Dulbecco’s PBS (1,000 × *g* for 30 sec. at RT), and eluted with 0.5 ml fractions of 1% formic acid. IgG-containing elution fractions were concentrated to dryness in a speed-vac, resuspended in ddH_2_O, combined, neutralized with 1M Tris / 3M NaOH, and prepared for liquid chromatography–tandem mass spectrometry (LC-MS/MS) as described previously (64, 65) with the modifications that (i) peptide separation using acetonitrile gradient was run for 120 min and (ii) data was collected on an Orbitrap Fusion (Thermo Fisher Scientific) operated at 120,000 resolution using HCD (higher-energy collisional dissociation) in topspeed mode with a 3 sec. cycle time. B cell receptor sequencing data was available from a previous study (30). Demultiplexing of sequence reads and the generation of consensus sequences for UMI groups were performed as outlined by Turchaninova *et al.* using software tools MIGEC (v1.2.9) and MiTools (v1.5) (66). Sequences with ≥2 reads were clustered into clonal lineages defined by 90% HCDR3 amino acid identity using USEARCH (67). LC-MS/MS search databases were prepared as previously described (64), using custom Python scripts (available upon request). MS searches, and MS data analyses were performed as previously described (64, 65), adjusting the stringency of the elution XIC:flowthrough XIC filter to 2:1.

### Data visualization and statistics

Flow cytometry data were analyzed and plotted using FlowJo (v10.7.1). Dot plots were generated using the package ggplot2 in RStudio (v1.4.1103) using R (v4.0.4). All other plots were generated in GraphPad Prism 9, which was also used for statistical analyses. The statistical test used for each analysis is indicated in the figure legends.

### Data availability statement

The BCR-seq data set analyzed in the current study is available in the NCBI SRA repository under accession numbers SAMN17497575-7.

## Conflict of interest

The authors declare that the research was conducted in the absence of any commercial or financial relationships that could be construed as a potential conflict of interest.

## Author contributions

EMB secured funding for the study, conceived the research question, and designed the study. SJG performed flow cytometry. SJG, KC, and SB produced recombinant antigens and monoclonal antibodies and contributed to other experiments. GB performed IFA and growth-inhibition assays. RAR, RG, and AEB performed merozoite ELISAs. AEB generated BCR-seq data. KCG and GCI performed plasma IgG proteomics. IS and BG provided clinical samples and data. SJG, SB, and EMB wrote the manuscript with input from all other co-authors.

## Funding

This work was supported by National Institutes of Health/National Institute of Allergy and Infectious Diseases (R01 AI153425 to E.M.B.). S.J.G and A.E.B. were supported by Graduate Research in Immunology Program training grant NIH T32 AI138944. R.A.R. was supported by Translational Science Training award TL1 TR002647. Data were generated in the Flow Cytometry Shared Resource Facility, which is supported by UT Health, NIH-NCI P30 CA054174-20 (CTRC at UT Health) and UL1 TR001120 (CTSA grant) and in the Genome Sequencing Facility, which is supported by UT Health San Antonio, NIH-NCI P30 CA054174 (Cancer Center at UT Health San Antonio), NIH Shared Instrument grant 1S10OD021805-01 (S10 grant), and CPRIT Core Facility Award (RP160732).

## Abbreviations

BCR: B cell receptor
HCDR3: heavy chain complementarity determining region 3
Ig: immunoglobulin
MBC: memory B cell
mAb: monoclonal antibody

## Acknowledgements

We would like to thank Dr. Kevin O. Saunders (Duke Human Vaccine Institute) for sharing reagents and protocols for the expression of recombinant IgG using linear Ig heavy and light chain gene expression cassettes from Liao *et al*. The 3T3-msCD40L cell line was a kind gift from Dr. Mark Connors (National Institute of Allergy and Infectious Diseases). Plasmids encoding 3D7 MSP1-bio, BirA, and rat CD200 were a kind gift from Dr. Gavin Wright, (Wellcome Sanger Institute; Addgene plasmids # 47709, 32408, and 36152). The following reagents were obtained through BEI Resources, NIAID, NIH: *Plasmodium falciparum*, Strain 3D7, MRA-102, contributed by Dr. Daniel J. Carucci; and Monoclonal Antibody N4-1F6 Anti-*Plasmodium falciparum* Apical Membrane Antigen 1 (AMA1) (produced *in vitro*), MRA-481A, contributed by Dr. Carole A. Long.

## Notes

### Competing Interest Statement

The authors have declared no competing interest.

