## Supplementary Information for "A comparative analysis of memory B cell and antibody responses against *Plasmodium falciparum* merozoite surface protein 1 in children and adults from Uganda"

**Supplementary table 1: Characteristics of donors used for MSP1-specific B cell isolations**

| Children | | | | Adults | | |
|---|---|---|---|---|---|---|
| **Donor ID** | **Gender** | **Age** | | **Donor ID** | **Gender** | **Age** |
| 170 | Male | 5 | | 1[1] | Unknown | ≥16 |
| 104 | Female | 5 | | 2[1] | Unknown | ≥16 |
| 3278 | Female | 6 | | 3071 | Male | 40 |
| 3280 | Male | 5 | | 3197 | Female | 41 |

[1] Donors were anonymous blood donors and not participants of a cohort study. Demographics and medical history are therefore not available.

**Supplementary table 2: Characteristics of donors used for anti-MSP1 plasma IgM and IgG reactivity measurements**

| | | | | Adults | | |
|---|---|---|---|---|---|---|
| **Donor ID** | **Gender** | **Age** | | **Donor ID** | **Gender[1]** | **Age** |
| 3059 | Female | 7 | | 3010 | Female | 37 |
| 3095 | Male | 8 | | 3017 | Male | 54 |
| 3102 | Male | 6 | | 3029 | Female | 40 |
| 3119 | Male | 5 | | 3034 | Female | 36 |
| 3149 | Female | 5 | | 3055 | Female | 44 |
| 3182 | Male | 6 | | 3068 | Female | 36 |
| 3199 | Male | 7 | | 3071 | Male | 40 |
| 3203 | Female | 6 | | 3075 | Female | 30 |
| 3210 | Female | 4 | | 3082 | Female | 41 |
| 3278 | Female | 6 | | 3132 | Female | 41 |
| 3280 | Male | 5 | | 3143 | Female | 32 |
| 3296 | Female | 7 | | 3165 | Female | 38 |
| 3297 | Male | 5 | | 3176 | Female | 41 |
| 3309 | Female | 6 | | 3197 | Female | 40 |
| 3316 | Male | 6 | | 3201 | Female | 41 |
| 3318 | Male | 8 | | 3223 | Female | 22 |
| 3326 | Male | 6 | | 3247 | Female | 72 |
| 3340 | Female | 5 | | 3249 | Female | 30 |
| | | | | 3283 | Female | 61 |
| | | | | 3295 | Female | 55 |
| | | | | 3311 | Male | 29 |
| | | | | 3324 | Female | 49 |
| | | | | 3347 | Female | 44 |
| | | | | 3418 | Female | 47 |

[1] In this predominantly pediatric cohort, samples from adults are acquired from caregivers, who are often female.

**Supplementary table 3: Molecular characteristics of anti-MSP1 plasma IgG as determined by proteomics analysis**

| Abun-dance (%) | Size of B cell lineage[1] | B cell subset[2] | Isotype | V-gene (IGH) | J-gene (IGH) | Amino acid changes in V-gene (%) | HCDR3 length (amino acids) |
|---|---|---|---|---|---|---|---|
| 24.7 | 1 | cMBC | IGHG3 | V3-53 | J5 | 22 | 10 |
| 23.5 | 5 | cMBC | IGHG3 | V1-69 | J6 | 26 | 24 |
| 15.5 | 1 | cMBC | IGHG3 | V3-64 | J1 | 25 | 18 |
| 14.0 | 1 | cMBC | IGHA1 | V4-28 | J5 | 16 | 10 |
| 6.9[3] | 4 | cMBC | IGHG1 | V1-69 | J6 | 25 | 21 |
| 4.5 | 1 | NBC | IGHD | V1-3 | J5 | 0 | 15 |
| 3.1 | 1 | atMBC | IGHG3 | V1-24 | J4 | 3 | 19 |
| 2.9 | 1 | cMBC | IGHG3 | V3-23 | J4 | 19 | 20 |
| 2.5 | 1 | NBC | IGHM | V4-28 | J5 | 7 | 18 |
| 1.2 | 1 | cMBC | IGHG1 | V4-39 | J4 | 22 | 15 |
| 0.4 | 1 | NBC | IGHM | V4-59 | J3 | 0 | 8 |
| 0.4 | 1 | cMBC | IGHG3 | V1-3 | J3 | 25 | 16 |
| 0.2 | 1 | cMBC | IGHG3 | V4-34 | J5 | 20 | 18 |
| 0.1 | 1 | atMBC | IGHM | V3-33 | J4 | 2 | 11 |
| <0.1 | 2 | atMBC | IGHM | V3-30 | J4 | 13 | 20 |
| <0.1 | 1 | NBC | IGHM | V3-53 | J3 | 0 | 9 |
| <0.1 | 1 | NBC | IGHM | V3-23 | J5 | 0 | 16 |
| <0.1 | 1 | NBC | IGHD | V3-53 | J4 | 0 | 14 |

[1] Number of unique clonal sequences found in BCR-seq data set that belonged to the same clonal lineage as the plasma IgG detected by proteomics analysis.

[2] BCR-seq subset in which a match was found with the sequence of anti-MSP1 plasma IgG.

[3] The lineage to which mAb10 and mAb22 belong is highlighted in green.

cMBC, classical memory B cell; NBC, naïve B cell; atMBC, atypical MBC.

**Supplementary table 4: Antibodies used for flow cytometry**

| Antibody | Fluorophore | Clone | Company / catalog number |
|---|---|---|---|
| CD19 | BV421 | SJ25C1 | BioLegend / 363017 |
| CD20 | BV785 | 2H7 | BioLegend / 302355 |
| CD21 | PerCP-eF710 | HB5 | Thermo / 46021942 |
| CD27 | PE-Cy7 | O323 | Thermo / 25027941 |
| IgA | FITC | IS11-8E10 | Miltenyi / 130-099-107 |
| IgD | PE-Dazzle594 | IA6-2 | BioLegend / 348240 |
| IgG | FITC | G18-145 | BD / 560952 |
| IgM | BV711 | MHM-88 | BioLegend / 314540 |
| IgM | eFluor 450 | SA-DA4 | Thermo / 48999841 |

**Supplementary table 5: Primers used in the study**

| # | **Sequence (5'-3')**[1] |
|---|---|
| 7 | CGCCTGAGTTCCACGACACC |
| 24 | **CTGGGTTCCAGGTTCCACTGGTGAC**CAGGTGCAGCTGGTRCAGTCTGGG |
| 25 | **CTGGGTTCCAGGTTCCACTGGTGAC**CAGRGCACCTTGARGGAGTCTGGTCC |
| 26 | **CTGGGTTCCAGGTTCCACTGGTGAC**GAGGTKCAGCTGGTGGAGTCTGGG |
| 27 | **CTGGGTTCCAGGTTCCACTGGTGAC**CAGGTGCAGCTGCAGGAGTCGG |
| 28 | **CTGGGTTCCAGGTTCCACTGGTGAC**GARGTGCAGCTGGTGCAGTCTGGAG |
| 29 | **CTGGGTTCCAGGTTCCACTGGTGAC**CAGGTACAGCTGCAGCAGTCAGGTCC |
| 30 | GCTGTGCCCCCAGAGGTGCTCYTGGA |
| 31 | **CTGGGTTCCAGGTTCCACTGGTGAC**GACATCCAGWTGACCCAGTCTC |
| 32 | **CTGGGTTCCAGGTTCCACTGGTGAC**GATATTGTGATGACCCAGWCTCCAC |
| 33 | **CTGGGTTCCAGGTTCCACTGGTGAC**GAAATTGTGTTGACRCAGTCTCCA |
| 34 | **CTGGGTTCCAGGTTCCACTGGTGAC**GACATCGTGATGACCCAGTCTC |
| 35 | **CTGGGTTCCAGGTTCCACTGGTGAC**GAAACGACACTCACGCAGTCTC |
| 36 | **CTGGGTTCCAGGTTCCACTGGTGAC**GAAATTGTGCTGACWCAGTCTCCA |
| 37 | **CTGGGTTCCAGGTTCCACTGGTGAC**GACATTGTGCTGACCCAGTCT |
| 38 | GGGAAGATGAAGACAGATGGT |
| 39 | **CTGGGTTCCAGGTTCCACTGGTGAC**CAGTCTGTGYTGACKCAGCC |
| 40 | **CTGGGTTCCAGGTTCCACTGGTGAC**CAGTCTGCCCTGACTCAGCC |
| 41 | **CTGGGTTCCAGGTTCCACTGGTGAC**TCYTATGAGCTGACWCAGCCAC |
| 42 | **CTGGGTTCCAGGTTCCACTGGTGAC**TCTTCTGAGCTGACTCAGGACCC |
| 43 | **CTGGGTTCCAGGTTCCACTGGTGAC**CAGCYTGTGCTGACTCAATC |
| 44 | **CTGGGTTCCAGGTTCCACTGGTGAC**CTGCCTGTGCTGACTCAGC |
| 45 | **CTGGGTTCCAGGTTCCACTGGTGAC**CAGSCTGTGCTGACTCAGCC |
| 46 | **CTGGGTTCCAGGTTCCACTGGTGAC**AATTTTATGCTGACTCAGCCCCACT |
| 47 | **CTGGGTTCCAGGTTCCACTGGTGAC**CAGRCTGTGGTGACYCAGGAG |
| 48 | **CTGGGTTCCAGGTTCCACTGGTGAC**CAGGCAGGGCWGACTCAG |
| 49 | GGGYGGGAACAGAGTGACC |
| 50 | AGTAATCAATTACGGGGTCATTAGTTCATAG |
| 51 | TCCCCAGCATGCCTGCTATTGTCTTCCCAATC |
| 52 | TCCCCAGCATGCCTGCTATTGTC |
| 53 | ATCCACTAGTAACGGCCGCCAGTG |

| | |
|---|---|
| 54 | CTGGGTTCCAGGTTCCACTGGTGAC |
| 55 | CACCTCTGGGGGCACAGC |
| 56 | CGAACTGTGGCTGCACCATCTGTCTTCATC |
| 57 | TGCCCCCTCGGTCACTCTGTTCCCGCCC |
| 58 | CAGTCACGACGTTGTAAAACGACG |
| 67 | GCTGTGCTCTCGGAGGTGCTCCTGGA |
| 106 | CAGCAGTGAGTAGAACCGTATCCG |
| 108 | ATGGCGGGAAGATGAAGACAG |
| 109 | AGTGTGGCCTTGTTGGCTTG |
| 110 | 5-Me-isodC/iso-dGCAGCAGTGAGTAGAACCGTATCCGrGrGrG |
| 297 | CCGACGGGGAATTCTCACAG |
| 298 | GCTGTGCCCCCAGAGGTGGAATTCTCACAGGAGACGAGG |
| 469 | GTAGAGGCTTGATTTGGAGGT |
| 494 | TGCCTATGCCTTATTCATCCCTC |

[1] Sequences indicated in bold do not bind to template but provide complementarity to the promoter or constant regions during the overlapping PCR to generate linear expression cassettes.

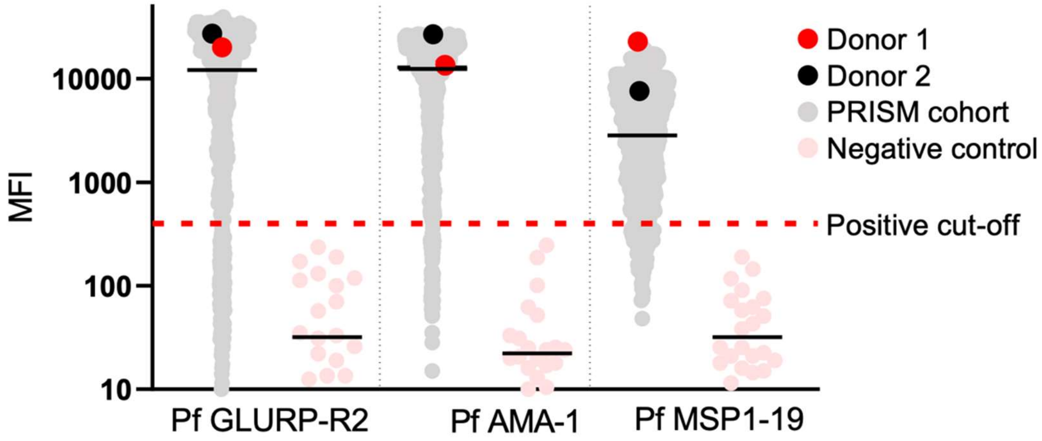


**Supplementary figure 1: Plasma antibody reactivity against *P. falciparum* antigens in malaria-experienced blood donors.** Antibody titers against three different parasite antigens were assessed by Luminex assay on plasma from malaria-experienced Ugandan blood donors (red and black). As a reference, antibody reactivity among participants of the PRISM cohort who live in a region of high *P. falciparum* transmission are also provided (gray). Black lines represent the median MFI of the PRISM cohort samples and the negative control samples (plasma from malaria-naïve European donors, pink). The mean of the negative control samples plus two standard deviations was used as the positive cutoff value. For all three antigens, the cutoff value was similar. The GLURP-R2 cutoff was the highest, and this value was chosen as the positive cutoff for the graph (dashed red line).

### Heavy chains

```
IGHV1-69*06 + IGHJ6*02

          <          FR1          >< CDR1 ><      FR2      >< CDR2 >
Germline  QVQLVQSGAEVKKPGSSVKVSCKASGGTFSSYAISWVRQAPGQGLEWMGGIIPIFGTA
mAb10     ..............................N.P....................V.A.T
mAb22     ............R..........I..DS.FNFP.....R..............V.A.T

          <              FR3                   ><         CDR3      ><   FR4   >
Germline  NYAQKFQGRVTITADKSTSTAYMELSSLRSEDTAVYYCAR           YYYYGMDVWGQGTTVTVSS
mAb10     .......D................V...K...........ALSRVRGVITH.............A.....
mAb22     D.SLR..D..A....Q..N......TN.KP..S.F.F.V.AVTRVRGVIVHH...AL....P........
```

### Light chains

```
IGKV3-11*01 + IGKJ4*01
          <          FR1           ><CDR1><      FR2      ><2>
Germline  EIVLTQSPATLSLSPGERATLSCRASQSVSSYLAWYQQKPGQAPRLLIYDAS
mAb10     .............................DN........A......I.....

          <              FR3                 >< CDR3 ><  FR4   >
Germline  NRATGIPARFSGSGSGTDFTLTISSLEPEDFAVYYCQQRSN LTFGGGTKVEIK
mAb10     K...........G...................L.......DE......N....R
```

```
IGLV2-14*03 + IGJL2*01
          <         FR1           ><  CDR1 ><      FR2      ><2>
Germline  QSALTQPASVSGSPGQSITISCTGTSSDVGGYNYVSWYQQHPGKAPKLMIYEVS
mAb22     ........................STG.I.Y................VIL.Q.N

          <              FR3                 >< CDR3  ><  FR4   >
Germline  NRPSGVSNRFSGSKSGNTASLTISGLQAEDEADYYCSSYTSS VLFGGGTKLTVL
mAb22     .......D...........Y.......DD.D...F.C.H...M............
```

**Supplementary figure 2: Heavy and light chain variable region sequences of mAb10 and mAb22.** The amino acid sequences of both monoclonal antibodies are aligned to their respective germline V-gene and J-gene sequences. Framework regions are indicated in blue and CDRs are indicated in yellow. Amino acid changes relative to the heavy chain variable region germline sequence that are shared between mAb10 and mAb22 are highlighted in orange.

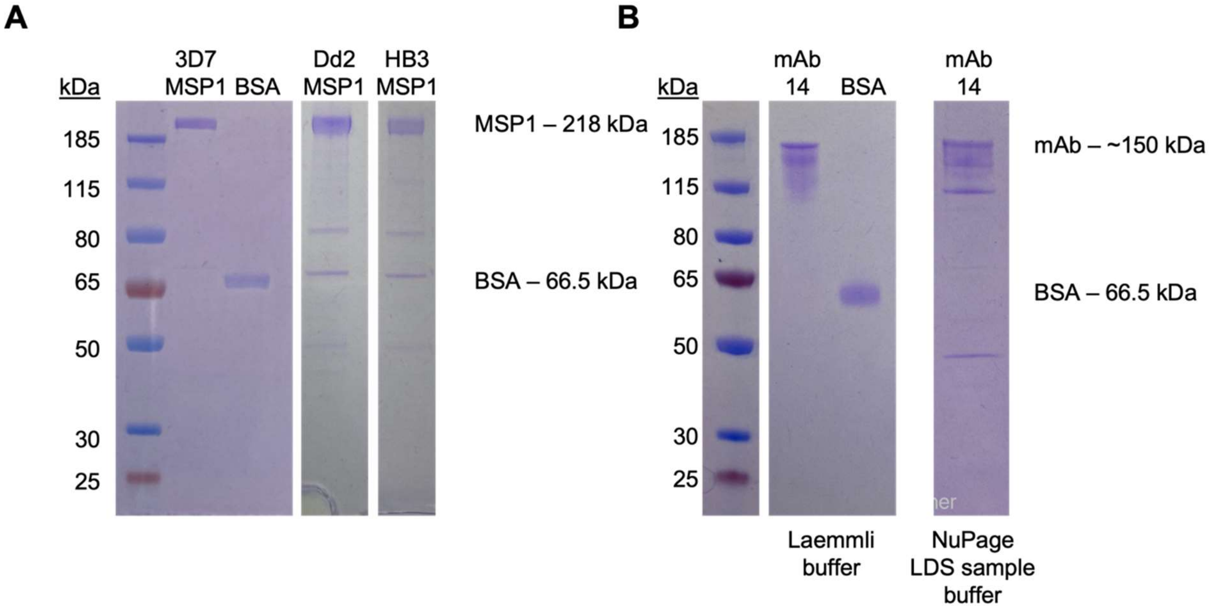


**Supplementary figure 3: Quality control of recombinant MSP1 and monoclonal antibodies by SDS-PAGE. A)** $MSP1_{3D7}$, BSA, $MSP1_{Dd2}$ and $MSP1_{HB3}$ (0.8 – 1.5 µg). **B)** mAb14 and BSA (0.5 - 1 µg). All proteins were loaded on gel in Laemmli buffer, with NuPage sample reducing agent for MSP1, but not mAbs. Loading mAbs in NuPage LDS sample buffer resulted in a pattern that resembled impartially formed immunoglobulins (mAb14 shown on the left).

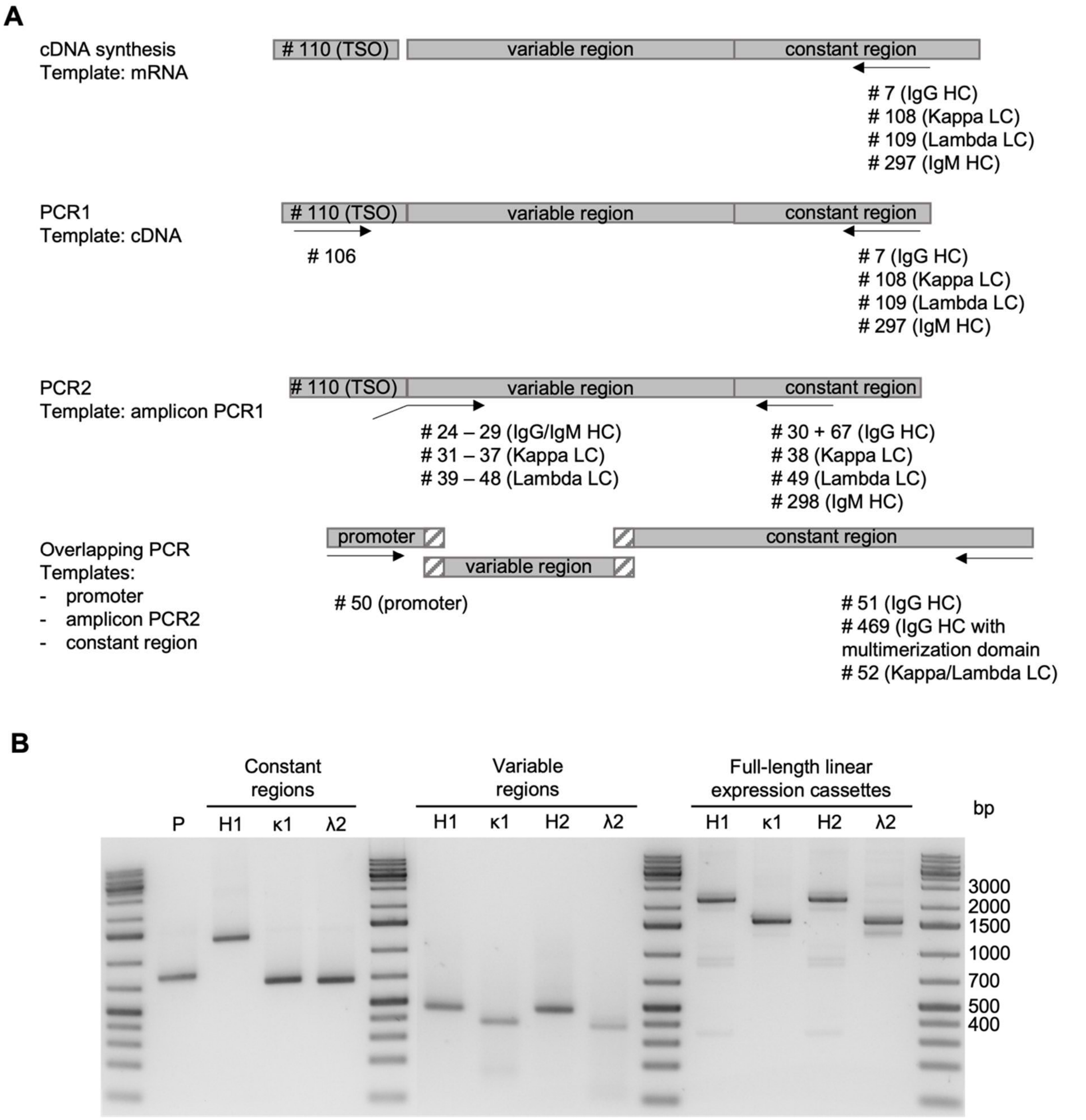


**Supplementary figure 4: Generation of linear antibody expression cassettes. A)** Schematic representation of reverse transcription and amplification of heavy and light chain variable regions from B cell mRNA. The sequences of primers indicated are listed in **Supplementary table 5**. **B)** DNA electrophoresis images showing the individual amplicons used in the overlapping PCR (P, promoter; H, heavy chain; κ, kappa light chain; λ, lambda light chain). The last four lanes show the full-length linear expression cassettes used for the expression of two recombinant monoclonal antibodies, one with a kappa light chain (H1 and κ1) and one with a lambda light chain (H2 and λ2).
